# Diffusible fraction of niche BMP ligand safeguards stem-cell differentiation

**DOI:** 10.1101/2022.09.13.507868

**Authors:** Sharif M. Ridwan, Autumn Twillie, Samaneh Poursaeid, Emma Kristine Beard, Muhammed Burak Bener, Matthew Antel, Ann E. Cowan, Shinya Matsuda, Mayu Inaba

**Affiliations:** Department of Cell Biology, University of Connecticut Health Center, Farmington, Connecticut, United States of America; Biozentrum, University of Basel, Basel, Switzerland; Richard D. Berlin Center for Cell Analysis and Modeling, University of Connecticut Health Center, Farmington, Connecticut, United States of America; Department of Molecular Biology and Biophysics, University of Connecticut Health Center, Farmington, Connecticut, United States of America

## Abstract

*Drosophila* male germline stem cells (GSCs) reside at the tip of the testis and surround a cluster of niche cells. It has been believed that the niche-derived Decapentaplegic (Dpp) has a role in maintaining stem cells in close proximity but has no role in the differentiating cells spaced one-cell layer away. However, the range of Dpp diffusion has never been tested. Here, using genetically encoded nanobodies called Morphotrap, we physically block Dpp diffusion without interfering with niche-stem cell signaling and find that diffusible fraction of Dpp is required to ensure differentiation of GSC daughter cells, opposite of its role in maintenance of GSC in the niche. Our work provides an example in which a soluble niche ligand induces opposed cellular responses in stem cells and in differentiating descendants so that the niche can tightly restrict its space. This may be a common mechanism to regulate tissue homeostasis.

**One sentence summary:** BMP ligand diffuses from the niche and has dual, and opposite roles on stem cells and differentiating daughter cells.

## Introduction

The stem cell niche was initially proposed to be a limited space in tissues or organs where tissue stem cells reside. Based on the phenomenon in which transplantation of hematopoietic stem cells is only successful when naïve stem cells are depleted, a niche is thought to provide a suitable environment for stem cells to self-renew (*1, 2*). At the same time, the niche environment should not foster the differentiation of descendant cells in order to ensure that the correct balance of self-renewal and differentiation is maintained (*2–4*). Although 40 years have passed since this niche concept was originally proposed (*1*), the mechanism of niche signal restriction is still poorly understood (*5*). This is partly because of the difficulty in studying stem cells in their *in vivo* context. Moreover, assessment of the dispersion of soluble molecules *in vivo* is challenging.

The *Drosophila* male germline stem cell system provides a model to study niche-stem cell interactions. The testicular niche, called the hub, is composed of post-mitotic hub cells. Each testis contains a single hub harboring 8-14 germline stem cells (GSCs) that directly attach to the hub (*6*). The division of a GSC is almost always asymmetric, via formation of a stereotypically oriented spindle, producing a new GSC and a gonialblast (GB), the differentiating daughter cell (*7*). After being displaced away from the hub, the GB enters 4 rounds of transit-amplifying divisions to form 2 to 16 cell spermatogonia (SGs). Then, 16-cell SGs become spermatocytes (SCs) and proceed to meiosis (Figure 1A-B) (*8*).

**Figure 1.**
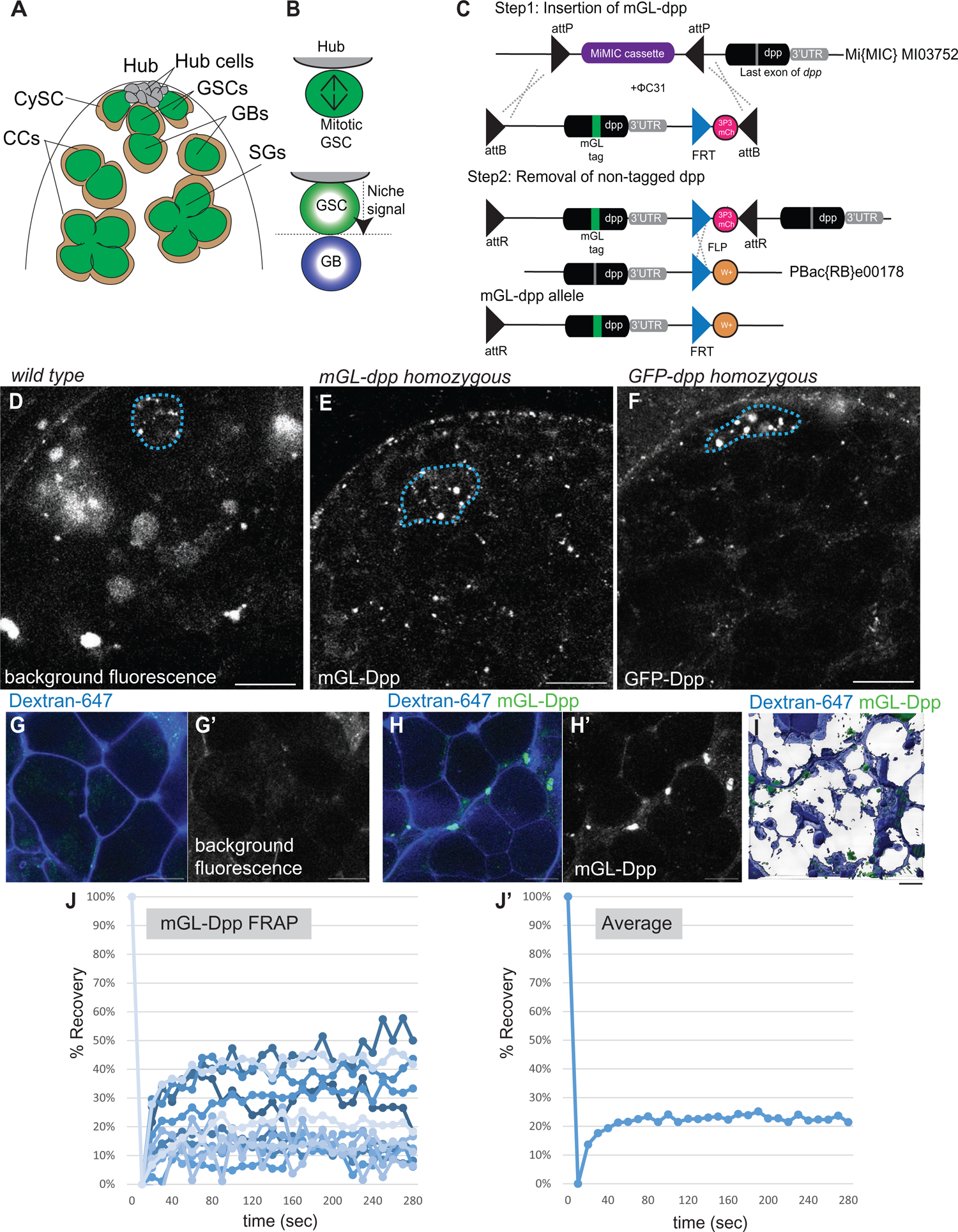
Diffusible fraction of Dpp is broadly observed in the testis. **A)** Anatomy of anterior area of *Drosophila* testis. Hub cells form a cluster and serve as the niche for germline stem cells (GSCs). Differentiating daughter cells or gonialblasts (GBs) undergo four rounds of incomplete division, called spermatogonia (SGs). Somatic cyst stem cells (CySCs) or cyst cells (CCs) are encapsulating developing germline. **B**) A schematic of asymmetric division (ACD) of GSCs. When the GSC divides, the mitotic spindle is always oriented perpendicularly towards hub-GSC interface (upper panel). As the result, GSC and GB are stereotypically positioned, one close to the hub and the other away from the hub (lower panel). Signal from the hub only activate juxtaposed daughter cell so that the two daughter cells can acquire distinct cell fates. **C**) A design of *mGreen Lantern (mGL)-dpp* allele. mGreen Lantern (mGL)-tagged *dpp* (last exon containing full-length *dpp* cording sequence after last processing site) with a 3xP3-mCherry cassette flanked by two inverted ΦC31 attB sites was replaced with gene trap cassette flanked by two inverted ΦC31 attP sites in MiMICMI03752. Then endogenous *dpp* last exon was removed by recombination between FRT sites of pBac(RB)e00178 and *mGL-dpp* cassette. *w*; dominant white marker. See details in *Method*. **D-F**) Representative images comparing testis tips isolated from wildtype (*yw,* **D**), homozygous *mGL-dpp* fly (**E**) or homozygous *GFP-dpp* knock-in fly (**F**) taken by the same microscope setting. **G-I**) Representative images of spermatogonial cysts after incubated with Alexa Fluor 647 conjugated dextran-dye. Wildtype (*yw*, **G**) *mGL-dpp* fly (**H**) and the 3D rendering (using Imaris) of a z-stack series of *mGL-dpp* (**I**). **J, J’**) Recovery curves of mGL-Dpp FRAP curves. % Recovery values (see Methods for calculation) from 13 trials are shown in **J**. Values from each trial are shown in different colors. **J’** shows average values of 13 trials. All scale bars represent 10 μm. Asterisks indicate approximate location of the hub. Live tissues were used for all images.

The Bone Morphogenetic Protein (BMP) ligand is often utilized in many stem cell niches in diverse systems (*9*). In the *Drosophila* testis, the BMP ligand, Decapentaplegic (Dpp) has emerged as a major ligand in the GSC niche together with a cytokine-like ligand, Unpaird (Upd) (*10–14*). In the testis, it has been hypothesized that these ligands activate signals in GSCs in close contact to the hub, and do not activate signals in GBs that are detached from the hub. However, the range of diffusion of these ligands and whether ligand diffusion beyond the niche space has a role is unknown.

We previously demonstrated that hub-derived Dpp is received by GSC-specific membrane protrusions, which we termed microtubule based (MT) - nanotubes, to efficiently activate downstream pathways within the GSC population (*15*). MT-nanotubes likely provide sufficient surface area along their length to allow the plasma membranes of GSCs and hub cells to closely contact one another for signaling (*15, 16*). This suggested the possibility that the Dpp signal is transmitted in a contact-dependent manner.

Furthermore, we serendipitously found that Dpp is almost freely diffusible (*17*), suggesting that Dpp could provide both contact-dependent and contact-independent signals. Besides the apparent contact-dependent signaling role of Dpp in the niche, we wondered what role the diffusing fraction of Dpp plays in the cells located outside of the niche.

In this study, we now directly address the function of the diffusing fraction of Dpp outside of the niche. We apply a previously established tool, Morphotrap, a genetically encoded nanobody that can trap secretory ligands on the plasma membrane of ligand-secreting cells (*18, 19*). Unexpectedly, we found that Dpp has distinct roles in differentiating germ cells from its actions in GSCs. In contrast to its GSC-specific function promoting the maintenance of GSCs, Dpp ensures differentiation of GB and SGs by blocking de-differentiation.

## Results

### Diffusible fraction of Dpp is broadly observed in the testis

We previously showed that overexpressed Dpp from the hub can diffuse outside of the niche (*17*). However, we were not able to successfully visualize a diffusing fraction of Dpp at endogenous protein levels to show that this behavior is physiological. In this study, we tackled this challenge by generating a fly line that expresses mGreen Lantern-tagged Dpp (*mGL-dpp*) from the endogenous locus, as described previously (*20*), so that we can monitor endogenous Dpp behavior (Figure 1C). Because mGL-Dpp signal was not distinguishable from background fluorescence in the heterozygous flies, we attempted to obtain homozygous flies. However, we found that the homozygous *mGL-dpp* allele was semi-lethal, hence we introduced a transgenic allele (*pPA dpp 8391/X*) containing genomic region of *dpp* expressed only in embryonic stage (*21*) to assist embryonic development. These rescued *mGL-dpp* homozygous flies were fully viable and able to reach adulthood with no phenotypes observed. These flies allowed us to successfully visualize endogenous Dpp expression and localization in the testis. In the hub, mGL-Dpp signal was seen as strong punctate pattern similar to the pattern previously shown in *mCherry-dpp* knock-in fly (Figure S1A, B) (*17*). In addition to the signal in the hub, the mGL-Dpp signal was seen throughout the testis towards the SC regions at levels above the background fluorescence (Figure 1D-F).

We noticed that mGL-Dpp localized in a pattern reminiscent of the extracellular space between cells throughout the anterior regions of testis and was not restricted to the niche (Figure 1E). To examine this further, we incubated testes isolated from *mGL-dpp* flies in media with freely diffusible fluorescent 10KDa dextran dye, which is of a similar size to Dpp protein (14.8kDa), for a short period of time (5-to-30 min). Previous work had shown that in such incubation with dextran, the dye penetrated in extracellular spaces between SG cysts and between germline and surrounding cyst cells (*22*). Indeed, we observed fluorescence accumulation in the testis tissue in a pattern matching previous description of extracellular space (Figure 1E). The dextran fluorescence appeared to penetrate between interconnected germ cells at various stages of SGs colocalizing with mGL-Dpp (Figure 1G-I). This suggested that mGL-Dpp localizes to extracellular space.

As the mGL-Dpp colocalizes to diffusible dextran dye outside the niche, we hypothesized that the mGL-Dpp signal resulted from long-range diffusion of the molecule throughout the tissue. To assess the mobility of the mGL-Dpp fraction in the tissue, we performed a fluorescence recovery after photobleaching (FRAP) analysis. After photobleaching, ∼20% of mGL-Dpp signal on average recovered within a few minutes (Figure 1J-J’), suggesting that there are two distinct fractions of the molecule present in the tissue: a mobile fraction (20% of the total) that is likely freely circulating, and an immobile fraction (80% of the total) that is likely bound to extracellular molecules/receptors or internalized by cells.

Taken together, these data indicate that Dpp likely freely diffuses from the niche and almost uniformly distributes throughout the testis. Supporting this data, we also observed similar distribution pattern of mScarlet-tagged Dpp (*mSC-dpp*) expressed from the endogenous locus allele (*22*) (Figure S1A, B). In contrast to these alleles, we observed the GFP-Dpp show almost undetectable fluorescent signal, and mCherry-Dpp shows highly concentrated signal within the hub as we reported previously (*17*) (Figure S1A, B). It should be noted that the GFP and mCherry tags were inserted into upstream of last processing site of Dpp, thus there is a possibility that non-tagged Dpp and/or free fluorescent proteins are present in the tissue. This may potentially cause the different pattern of fluorescent signal in the testis (Table S1).

### Perturbation of Dpp diffusion without affecting niche-GSC signal

Since differentiating cells are descendant of stem cells, it is difficult to determine the direct and specific effect of niche ligands on differentiating cells. In fact, while Dpp function within the niche is well-characterized, the role of a potentially diffusible Dpp fraction outside of the niche is completely unknown. In order to assess the function of the diffusible fraction of Dpp, we sought to specifically disturb only diffusing fraction of Dpp, without affecting the niche-GSC Dpp signal. To achieve this, we utilized the Morphotrap (MT), a genetically encoded tool consisting of a fusion protein between a transmembrane domain and a nanobody that recognizes green fluorescent protein (GFP) including its variants, and acts as a synthetic receptor for GFP-tagged proteins (*18, 19*). We used two versions of MT, each expressing a fusion protein of anti-GFP nanobody and one of two different transmembrane proteins, Nrv1 or mCD8 (Figure 2A).

**Figure 2.**
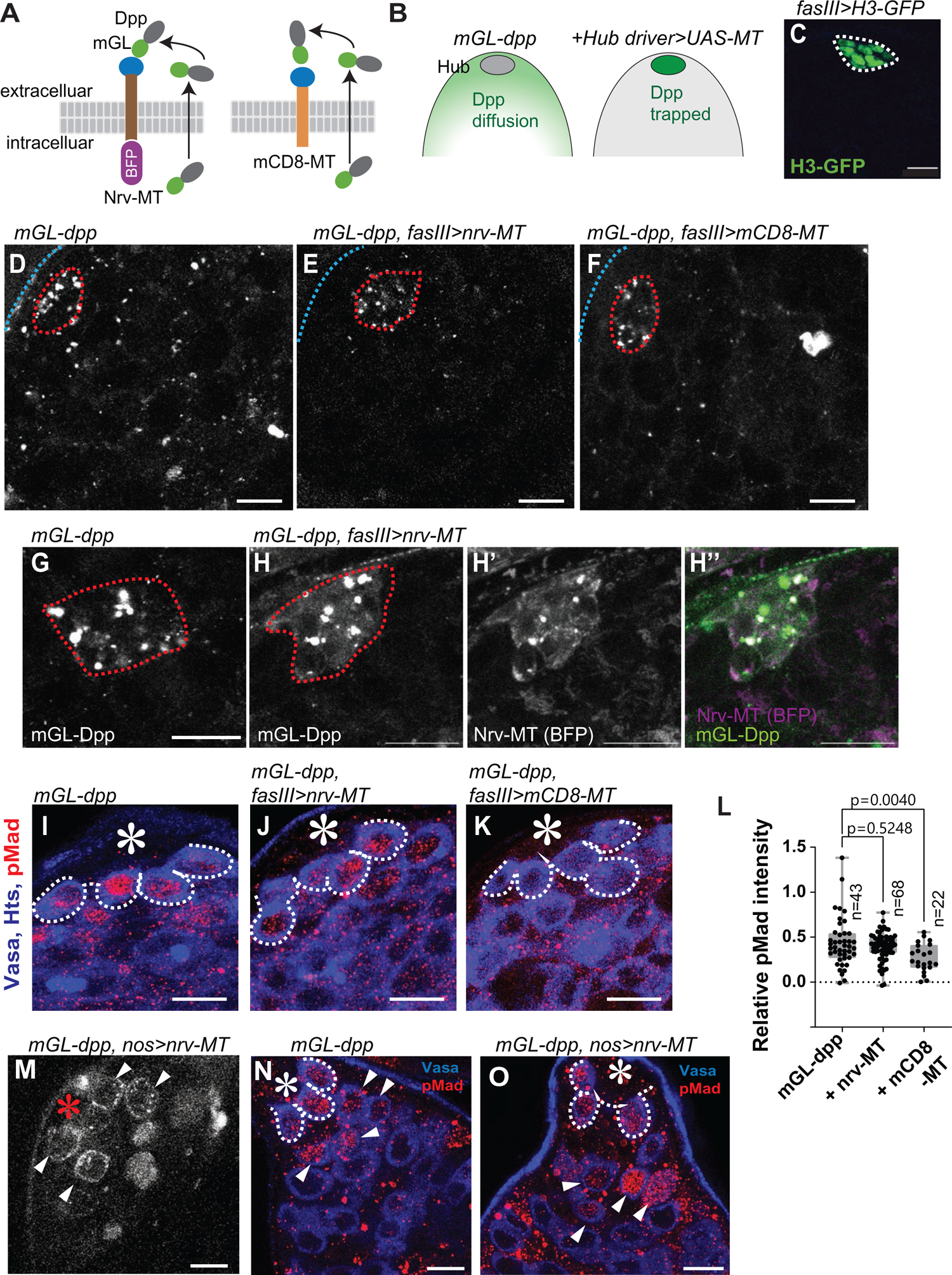
Perturbation of Dpp diffusion without affecting niche-GSC signal. **A)** Schematics of the design to trap Dpp on the surface of Dpp producing cells using Morphotrap (MT), the genetically encoded synthetic receptor for GFP-tagged proteins. The nanobody, vhhGFP4 (blue circle), that specifically binds to GFP, is fused to transmembrane domains of either mouse CD8 (mCD8-MT) or Nrv1 basolateral scaffold protein (Nrv-MT). **B**) Expected outcome of hub-driven expression of morphotrap in the background of *mGL-dpp* homozygous fly rescued by *pPA dpp 8391/X*. Diffusing fraction of Dpp (left panel) will be trapped on the hub cell surface and no diffusing Dpp will be observed (right panel). **C**) Representative images of Histone H3-GFP expressed under the hub-specific driver, fasIIIGal4. **D, F**) Representative images of testis tip of homozygous *mGL-dpp* fly rescued by *pPA dpp 8391/X* without (**D**) or with (**E**) fasIIIGal4 driven Nrv-MT expression or with (**F**) fasIIIGal4 driven mCD8-MT expression. Hub is encircled by red broken lines. **G-H**) Magnification of hub area from homozygous *mGL-dpp* fly rescued by *pPA dpp 8391/X* without (**G**) or with (**H**) fasIIIGal4 driven Nrv-MT expression. **I-K**) Representative images of pMad staining of GSCs after trapping Dpp using indicated Morphotrap lines. The fasIIIGal4 driver was used. White broken lines encircle GSCs. **L)** Quantification of pMad intensity in GSCs (relative to CCs) of fasIIIGal4 driven Nrv-MT or mCD8-MT expressing testes in *mGL-dpp* homozygous background rescued by *pPA dpp 8391/X*. P-values were calculated by Dunnett’s multiple comparisons tests. “n” indicates the number of scored GSCs. Box plots show 25–75% (box), minimum to maximum (whiskers) with all data points. **B) M)** Representative images of live testis tip of homozygous *mGL-dpp* fly rescued by *pPA dpp 8391/X* expressing Nrv-MT under the germline specific driver, nosGal4. Trapped mGL-Dpp signal is seen on the surface of early germ cells (white arrowheads). **N, O**) pMad staining shows emerging pMad positive germ cells (arrowheads in **O**) outside of the niche in the same genotype used in **M**. pMad positive germ cells are normally only seen in GSCs and immediate descendants around the hub (arrowheads in **N**). White broken lines encircle GSCs. All scale bars represent 10 μm. Asterisks indicate approximate location of the hub. Live tissues were used for **C-H, M** and fixed samples were used for **I-K, N, O**.

Nrv-MT consists of the Nrv1 protein scaffold and localizes to the basolateral compartment of the *Drosophila* wing disc (*19*). mCD8-MT consists of the membrane protein mCD8 and localizes throughout the entire plasma membrane (*19*). In order to trap Dpp on the surface of niche cells, we utilized the hub driver fasIIIGal4, which drives expression specifically in the hub cells that make up the germline stem cell niche (Figure 2B, C). By expressing MT under control of the fasIIIGal4 driver in the *mGL-dpp* homozygous background, we reasoned that we could remove all circulating fraction of Dpp, including fractions secreted from the hub cells or from any other cell types, to prevent its effect outside of the niche, and tether them on hub cell membranes to keep it to signal to stem cells via a contact dependent manner (Figure 2B).

Indeed, we found that expression of both Nrv-MT and mCD8-MT under the fasIIIGal4 driver eliminated mGL-Dpp signal throughout the testis (Figure 2D-F), indicating that both MTs can efficiently trap mGL-tagged Dpp. In contrast, we observed trapped mGL-Dpp on the surface of hub cells (Figure2G-H). Interestingly, although we were expecting to see strong signal on the surface of hub cells, the signal was not as high as reflecting equivalent amount to all diffusing fraction of mGL-Dpp. Therefore, we wondered if trapped mGL-Dpp may be constantly internalized and degraded in hub cells. If so, we should see increased level of mGL signal in hub lysosomes when lysosome digestion is perturbed by chloroquine (CQ) treatment, a drug that inhibits lysosomal enzymes. Indeed, the Dpp-mGL signal in hub lysosomes after 4-hour chloroquine treatment showed increase in Dpp-mGL in hub cells of MT expressing testes (Figure S2A-C), indicating that trapped Dpp may be constantly degraded in hub cells.

Next, to examine whether MT expression specifically perturbs diffusible fraction of Dpp and keep hub-GSC signal intact, we stained the testis for phosphorylated Mad (mothers against dpp) protein, a readout of Dpp signal activation. In a normal testis, levels of phosphorylated Mad (pMad) are high in GSCs, and it immediately becomes lower in GB/SGs. We found that pMad intensity was reduced in GSCs in mGL-Dpp, fasIII>mCD8-MT testes compared to control testes (Figure 2G-J). In comparison, mGL-Dpp, fasIII>Nrv-MT showed similar pMad intensities in the GSCs as compared to control sample (Figure 2G-J), indicating that hub-GSC signal was unaffected. Note that trapping Dpp did not perturb pMad in somatic cyst cells in both samples (Figure S1D-G), which suggests that pMad in cyst cells is not activated by circulating Dpp and serves as a reliable internal control for quantifying relative pMad intensity in germ cells.

In contrast to the MT expression in hub cells, in which trapped mGL-Dpp is internalized and degraded, expression of Nrv-MT using the germline driver nosGal4 (mGL-Dpp, nos>Nrv-MT) clearly visualized mGL-Dpp trapping along the membranes of germ cells outside of the niche (Figure 2K, L) and hyper-activation of signaling as indicated by elevated pMad staining (Figure 2M, N). These data confirm that a fraction of Dpp is diffusible and trappable by the MT method, and that trapped Dpp can still signal to receptors present on the plasma membrane of the cells.

It should be noted that knock-down of mGL-tagged Dpp in the hub (*pPA dpp 8391/X*, *fasIII>GFP-IR, mGL-dpp/mGL-dpp*) significantly reduced pMad signal in GSCs, confirming that rescuing transgene (*pPA dpp 8391/X*) does not contribute to the expression in the hub (Figure S2H). Based on these results, we concluded that fasIII>Nrv-MT expression in the *mGL-dpp* homozygous (rescued with *pPA dpp 8391/X*) background was the best tool to be used to assess the function (s) of diffusible fraction of Dpp outside of the niche, without disrupting hub-GSC Dpp signaling.

### The diffusible fraction of Dpp prevents de-differentiation

In the *Drosophila* testis, GSCs almost exclusively divide asymmetrically to produce one GSC and one GB (asymmetric outcome, Figure 3A) (*7, 23, 24*). However, symmetric event can also occur (symmetric outcome, Figure 3A) (*23, 24*) via two mechanisms: 1) spindle misorientation, where the mitotic spindle orients parallel to the hub-GSC interface, resulting in two GSCs (Figure 3A-17) (*7*), and 2) de-differentiation, where a differentiating GB or SG physically relocates back to the niche and reverts to a GSC identity (Figure 3A-17) (*25*). Although a recent study suggested that the de-differentiation from the Bam-positive lineage (4-16-cell SGs) is required for maintenance of stem cell number only in challenging conditions (*26*), it was observed constantly from the earlier lineage (GB to 2-cell SGs) even in physiological conditions (*24*), indicating that de-differentiation is a critical mechanism to maintain stem cell numbers. At the same time, excessive de-differentiation is hypothesized to be a cause of tumorigenesis (*27*).

**Figure 3.**
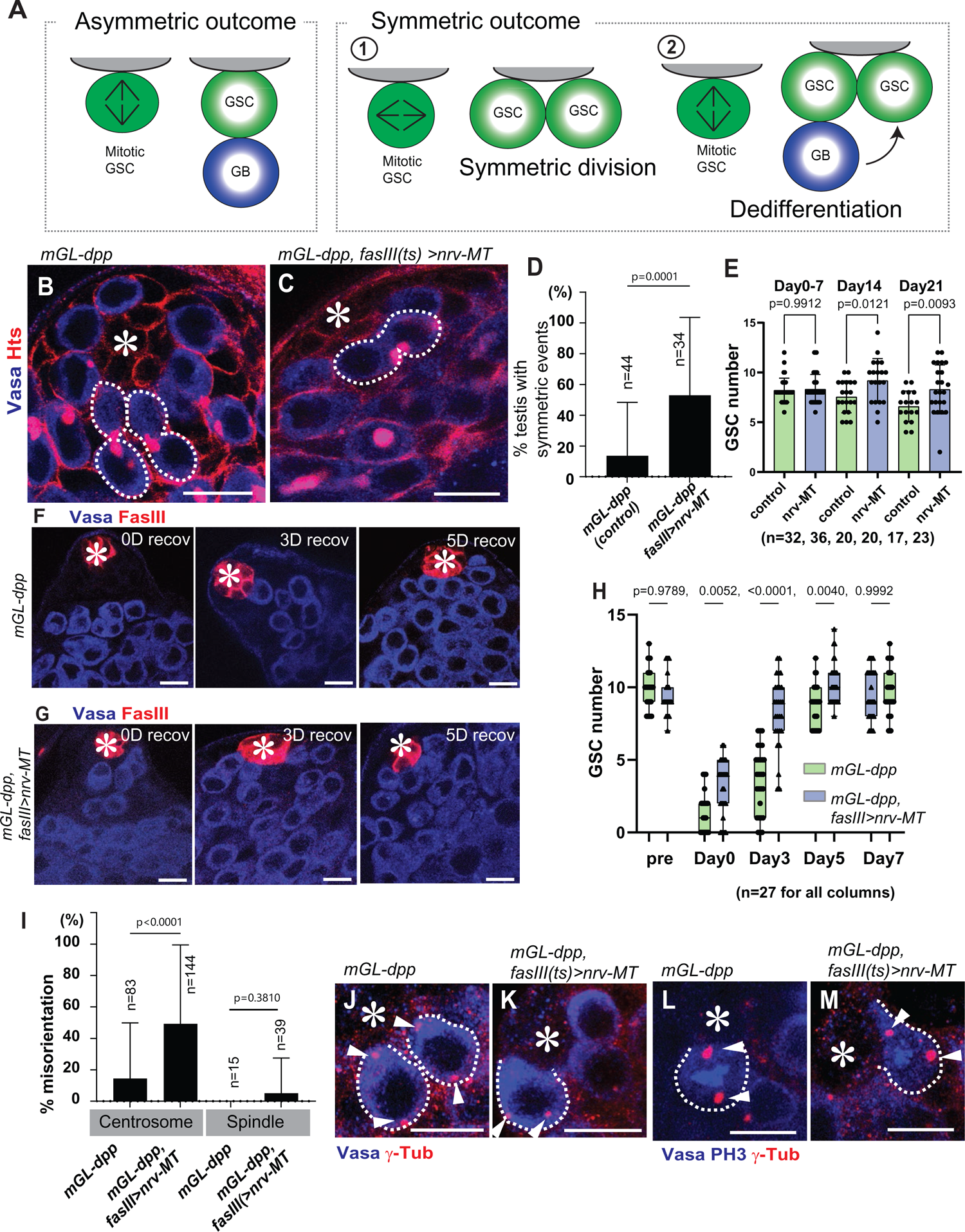
The diffusing fraction of Dpp prevents de-differentiation outside the niche. **A)** Asymmetric and symmetric outcomes of GSC division. Symmetric outcome is defined as the case in which two daughter cells of a GSC division are both placed near the hub, resulting in production of two GSCs. It occurs as the consequence of either “symmetric division” (1) or “de-differentiation” (2) (see details in main text). **B, C**) Representative images of testis tip without (**B**) or with (**C**) trapping Dpp. Broken lines indicate asymmetric events in **B** and a symmetric event in **C**. **D**) Frequency of testes showing any symmetric events without or with trapping Dpp. **E**) Changes in GSC number during aging without or with trapping Dpp. **F, G**) Representative images of testis tip after depletion of GSC by expressing Bam at 0-day recovery (0D recov: immediately after 6-time heat shock treatment) and after 3-day and 5-day recovery time points (3D recov, 5D recov) in room temperature culture without (**F**) or with (**G**) trapping Dpp. **H**) Changes in GSC number during recovery from forced differentiation of GSCs without or with trapping Dpp. Pre: pre-heat shock. **I**) Percentages of misoriented centrosome and spindle in GSCs without or with trapping Dpp. **J-M**) Representative images of centrosomes in interphase cells (**J, K**) and mitotic cells (**L, M**) of GSCs without (**J, L**) or with (**K, M**) trapping Dpp. For trapping Dpp in this figure, Nrv-MT was expressed under the control of fasIIIGal4 driver in *mGL-dpp* homozygous background rescued by *pPA dpp 8391/X*. no Gal4 flies were used for control. Fixed samples were used for all images and graphs. All scale bars represent 10 μm. Asterisks indicate approximate location of the hub. “n” indicates the number of scored testes in **D**, **E** and **H**, or scored GSCs in **I**. Box plots show 25–75% (box), minimum to maximum (whiskers) with all data points. Other graphs show means and standard deviations. The p-value was calculated by student-t-test for **D** and **I** and by Šídák’s multiple comparisons tests for **E** and **H** and provided on each graph.

By scoring the orientation of cells still interconnected by the fusome, a germline-specific organelle that branches throughout germ cells during division, we can estimate the frequency of symmetric events in the niche (*24, 25, 28*). We noticed that mGL-Dpp, fasIII>Nrv-MT testes showed a significantly higher frequency of symmetric events than the control (Figure 3B-D), suggesting that preventing Dpp diffusion results in more symmetric outcomes of GSC division. In mGL-Dpp, fasIII>Nrv-MT testes, the number of GSCs at the hub were slightly higher at timepoints of day 14 and 21 post-eclosion (Figure 3E), suggesting the possibility that preventing Dpp diffusion may cause excess de-differentiation.

In the *Drosophila* testis, Bag of marbles (Bam), a translational repressor, is expressed after a germ cell exits the GSC state and is sufficient for promoting differentiation (*25*). Using heat-shock inducible expression of Bam in GSCs, we can artificially induce differentiation of GSCs, resulting in depletion of all GSCs from the niche. After the flies are recovered in normal temperature, the niche is replenished through de-differentiation (*25*). By introducing *hs-bam* transgene in the *mGL-dpp* homozygous background rescued by *pPA dpp 8391/X* with or without fasIII>Nrv-MT expression, we assessed the function of the diffusible Dpp fraction on de-differentiation. Strikingly, induction of GSC differentiation in mGL-Dpp, fasIII>Nrv-MT flies resulted in a significantly faster recovery of GSCs in the niche as compared to flies without fasIII>Nrv-MT expression (Figure3F, H). Moreover, Dpp trapping using an alternative driver/knock-in combination (*dppGal4>nrv-MT, GFP-dpp* knock-in homozygous background, note that this line is homozygous viable and no transgenic was used for the rescue) showed similar effect on de-differentiation (Figure S3), confirming that the trapping of diffusible Dpp is the cause of observed phenotype.

These data suggests that diffusible Dpp plays a role in preventing de-differentiation in differentiating GBs and SGs. Since the Nrv-MT only affected differentiating cells located away from the niche, but not GSCs within the niche, we speculated that increased de-differentiation rather than increased spindle misorientation may be responsible for observed high frequency of symmetric events. To further rule out the possibility that defects in spindle orientation in the GSCs contributes to the effects, we assessed the spindle orientations of GSCs in the mGL-Dpp, fasIII>Nrv-MT flies. We found that GSCs showed correctly oriented spindles (Figure 3I-M), in support of our conclusions that that observed symmetric outcome are the result of excess de-differentiation and not spindle misorientation. We do observe that although the spindles were correctly oriented, centrosomes of GSCs in mGL-Dpp, fasIII>Nrv-MT flies were significantly more misoriented (Figure 3I-M), but this is likely a secondary effect of a higher frequency of de-differentiation as de-differentiated GSCs are reported to have higher instances of centrosome misorientation (*29*).

### Dpp acts through its canonical pathway in GSCs and in differentiating germ cells

We next asked if Dpp acts within the same signaling pathway in differentiating germ cells as it does in GSCs. Dpp is known to bind to its receptor Thickveins (Tkv) on GSCs and activate Tkv-mediated signaling to maintain GSC identity (*10, 11*). Knock-down of Tkv by expression of shRNA under the control of the germline driver nosGal4 results in a depletion of GSCs from the niche (Figure 4A-B), demonstrating the indispensability of this pathway to GSC maintenance consistent with previous reports (*10, 11*). To determine if Tkv is the receptor for diffusible Dpp for germ cells outside of the niche, we knocked down Tkv exclusively in differentiating germ cells using the driver bamGal4.

**Figure 4.**
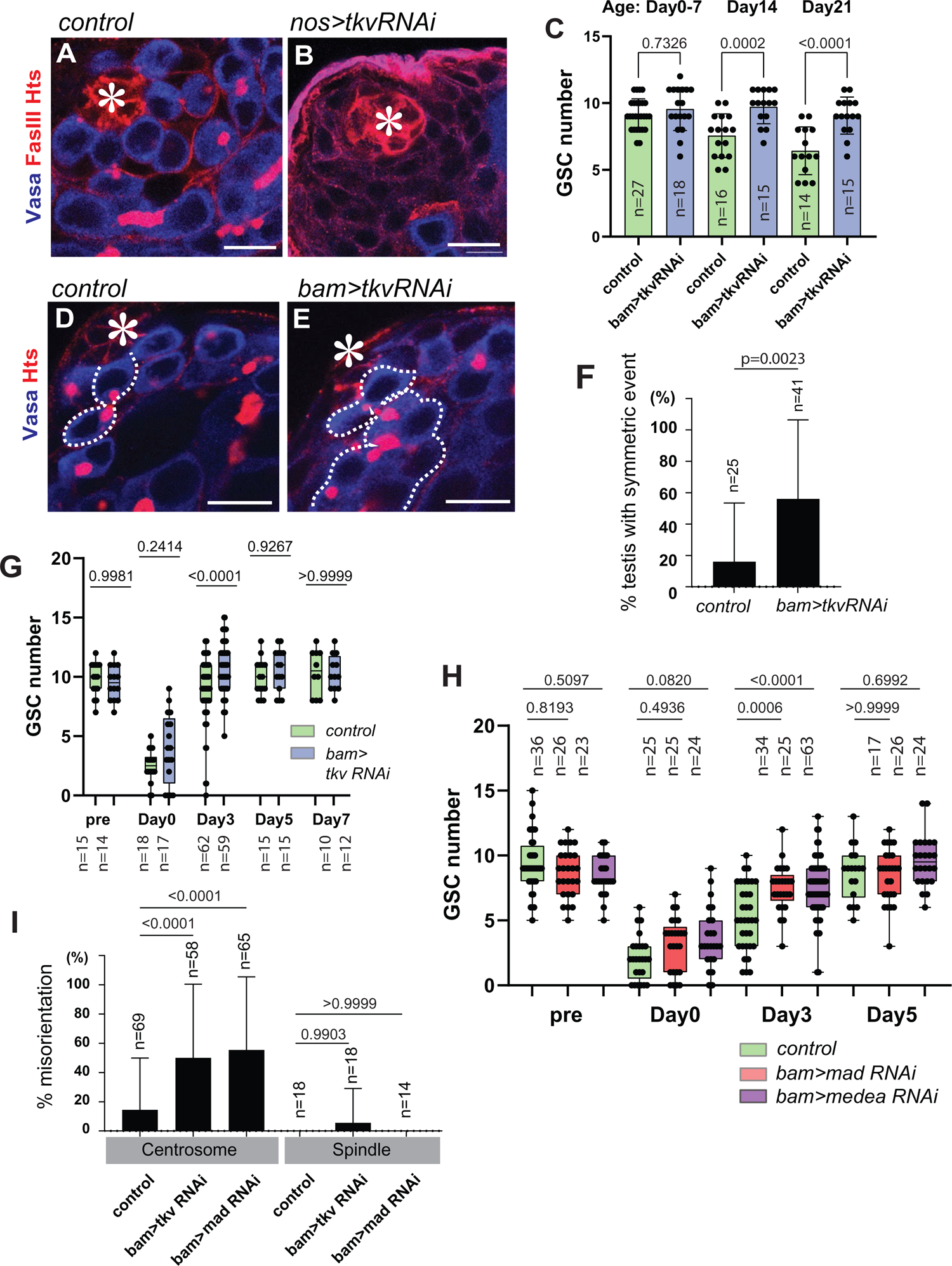
Dpp acts through its canonical pathway in GSCs and in differentiating germ cells. **A, B**) Representative images of testis tip without (**A**) or with (**B**) expression of shRNA against tkv (*tkv RNAi*) under the nosGal4 driver. Tkv RNAi shows the hub without harboring any Vasa positive germ cells. **C**) Changes in GSC number during aging without or with *tkv RNAi* expression under the bamGal4 driver. **D, E**) Representative images of testis tip without (**D**) or with (**E**) *tkv RNAi* under the bamGal4 driver. Broken lines indicate symmetric events. **F**) Frequency of testes showing any symmetric events without or with *bam>tkv RNAi*. **H**) Changes in GSC number during recovery from forced differentiation of GSCs without or with bam>Tkv RNAi (**G**), Mad RNAi, Medea RNAi (**H**). **I**) Percentages of misoriented centrosome and spindle in GSCs of indicated genotypes. All scale bars represent 10 μm. Asterisks indicate approximate location of the hub. Fixed samples were used for all images and graphs. “n” indicates the number of scored testes in **C**, **F**, **G** and **H**. Box plots show 25–75% (box), minimum to maximum (whiskers) with all data points. Other graphs show means and standard deviations. The p-value was calculated by student-t-test for **F** and by Šídák’s multiple comparisons tests for **C, G, H, I** and provided on each graph.

Intriguingly, we observed a higher number of GSCs per niche in bam>Tkv RNAi testes as flies aged (Figure 4C), similar to what was observed in mGL-Dpp, fasIII>Nrv-MT flies (Figure 3E). Moreover, bam>Tkv RNAi testes also exhibit a higher frequency of symmetric events (Figure 4D-F), recapitulating the phenotype of mGL-Dpp, fasIII>Nrv-MT flies and suggesting that Tkv-mediated signaling in differentiating germ cells may similarly result in higher instances of de-differentiation. Indeed, analysis of hs-bam with bam>Tkv RNAi show a significantly faster recovery of GSCs than control after heat-shock mediated depletion of GSCs (Figure 4G), indicating Tkv-mediated signaling in differentiating cells impedes de-differentiation.

To further confirm that the observed faster recovery is surely caused by accelerated de-differentiation, we conducted long-term live imaging of bam>Tkv RNAi flies to measure the frequency of de-differentiation events (Figure S4A-C, supplemental video S1, 2). We monitored the division of GSCs expressing mCherry-Vasa for 16 hours. Strikingly, we found dramatic increase of frequency of de-differentiation events in Tkv RNAi expressing testes, indicating that diffusible fraction of Dpp prevents de-differentiation.

Next, we knocked-down Mad, the downstream effector of Tkv-signaling, and Medea, the partner of Mad, using bamGal4 mediated shRNA expression in hs-bam flies. We found that both RNAi conditions show a significantly faster recovery of GSCs after heat-shock mediated GSC depletion, indicating the Tkv-Mad/Medea pathway that is responsible for maintenance of GSCs, is also responsible for preventing de-differentiation (Figure 4H).

As was the case of mGL-Dpp, fasIII>Nrv-MT flies, spindles were not misoriented in *bam>tkv RNAi*, *bam>mad RNAi* or *bam>medea RNAi* genotypes (Figure 4I), again suggesting that de-differentiation, and not spindle misorientation, is responsible for the increase in symmetric events. Centrosomes for these genotypes did exhibit misorientation but as noted above, this is a phenomenon frequently seen in GSCs as a consequence of de-differentiation.

These results strongly suggest that both the contact dependent and independent signal from Dpp (inside and outside of the niche) use same signaling pathway to achieve distinct signaling outcomes.

### Gbb, Sax and Punt are required for preventing de-differentiation

In addition to Dpp, it has been reported that another BMP ligand, Glass bottom boat (Gbb), the Drosophila BMP7 ortholog, is also required for GSC maintenance in the testis (*11*). Therefore, we decided to investigate the function of Gbb in GSC maintenance and differentiation. First to understand expression pattern of Dpp and Gbb, we conducted anti-HA staining of fully functional HA-tagged Dpp and Gbb (*30, 31*) to detect intracellular fraction of Dpp and Gbb. In a past report, Dpp was shown to be expressed both in hub cells and CySCs by RT-PCR (*11*).

However, our staining showed that Dpp protein exclusively in hub cells, whereas Gbb is expressed not only in hub cells, but also in CySCs and CCs (Figure 5A-C). Dpp-mGL also showed similar intracellular pattern when extracellular mGL-Dpp signal was removed by fixation (Figure 5D-F). These data suggest that Dpp and Gbb have distinct expression pattern at least in their protein level.

**Figure 5.**
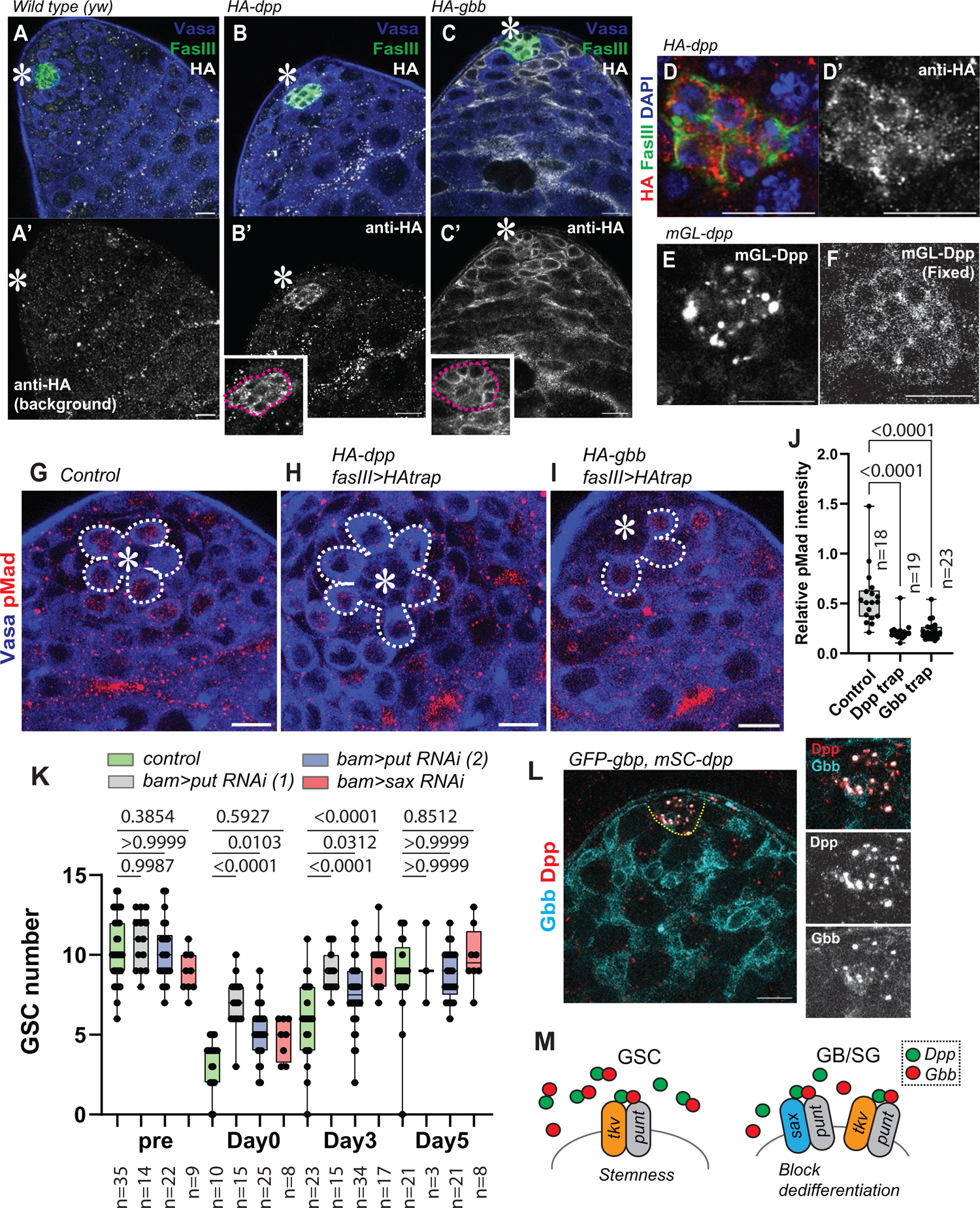
Gbb, Sax and Punt are required for preventing de-differentiation. **A-C)** Representative images of HA staining of indicated genotypes. Homozygous *HA-dpp* and *HA-gbb* flies were used. **A’-C’** show anti-HA channel. Right panels show magnified areas of the hub. **D**) A representative image of the hub of *HA-dpp* testis. **D’** shows anti-HA channel. **E-F**) Comparison of mGL-Dpp signal before (**E**) and after (**F**) fixation. **G-I**) Representative images of pMad staining of GSCs without (**G**), after trapping Dpp (**H**) or Gbb (**I**) by expressing HAtrap under the FasIIIGal4 driver. GSCs are encircled by white dotted lines. **J**) Quantification of pMad intensity in GSCs (relative to CCs) of fasIIIGal4 driven HAtrap in the background of *HA-dpp* homozygous fly or *HA-gbb* homozygous fly. P-values were calculated by Dunnett’s multiple comparisons tests. **K**) Changes in GSC number during recovery from forced differentiation of GSCs without or with bam> punt RNAi (1) TRiP.GLV21066, punt RNAi (2) TRiP.HMS01944 or sax RNAi:TRiP.HMJ02118. P-values were calculated by Šídák’s multiple comparisons test. **L**) A representative image of colocalization of GFP-Gbb and mSC-Dpp in the hub. Images were taken using live testes. **M**) Model. Dpp and Gbb may form heterodimer and secreted from hub cells induce distinct signaling outcomes in GSC and in GB/SG. Tkv functions both in GSCs and GB/SGs, whereas Sax predominantly functions in GB/SGs. All scale bars represent 10 μm. Asterisks indicate approximate location of the hub. Box plots show 25–75% (box), minimum to maximum (whiskers) with all data points. “n” indicates the number of scored GSCs in **J**, testes in **K**.

Next, we conducted trapped HA-tagged Dpp or Gbb using HA-trap (*30–32*). Trapping HA-Dpp showed significant reduction of pMad in GSCs similar to trapping mGL-Dpp by mCD8-MT (Figure 5G, H and J). Interestingly, trapping Gbb-HA in hub cells was enough to reduce pMad in GSCs, despite its expression in CySCs and CCs. In contrast, knock-down of Gbb in CySCs and CCs under the c587-Gal4^ts^ driver did not affect signal in GSCs (Note that this condition affected pMad signal in CCs, Figure S5A, B). These data suggest that hub-derived Gbb but not CySC/CC-derived Gbb can signal to GSCs (Figure 5I, J, Figure S5A, B).

Since trapping of HA-tag on Gbb-HA in hub cells showed significant reduction of pMad in GSCs, we could not use HA-trap to test the function of Gbb specifically outside of the niche. Therefore, we tested involvement of a type I receptor, Saxophone (Sax), which preferentially binds to Gbb (*33*), and Punt, a common type II receptor for Dpp and Gbb (*32*). Knock-down of Sax or Punt under bamGal4 driver both showed acceleration of dedifferentiation, similar to the phenotype of mGL-Dpp trap and Tkv RNAi, indicating that both Dpp and Gbb outside of the niche are required for preventing dedifferentiation (Figure 5K).

Interestingly, knock-down of Punt but not Sax under the control of the germline driver nosGal4 resulted in an immediate depletion of GSCs from the niche (Figure S5C-F), demonstrating the requirement of Sax to GSC maintenance is minor. Moreover, pMad was strongly positive in Sax RNAi GSCs (Figure S5C, D), suggesting that Sax mainly works outside of the niche.

Based on these results, we propose that both ligands, Dpp and Gbb, may have critical effect both on GSC maintenance and prevention of dedifferentiation. Despite Dpp shows uniform distribution over the entire tissue and Gbb has broad expression pattern, we observed co-localization of Dpp (mScarlet-Dpp) and Gbb (GFP-Gbb) predominantly within the hub, where they may contribute to the formation of sharply graded signaling outcomes in the cells around the niche (Figure 5L, M).” A recent report has demonstrated that the heterodimers of Dpp-Gbb form within a cell prior to secretion and that it triggers strong signaling (*31*). It is possible that Dpp/Gbb heterodimer may form specifically in the hub and contribute to graded response of cells.

### Diffusible fraction of Dpp activates Bam expression in differentiating germ cells to prevent de-differentiation

It is known that Dpp signal suppresses expression of the bam gene in female GSCs where Bam is necessary and sufficient for differentiation and its suppression in stem cells is essential to maintain their undifferentiated states in female GSCs (*34, 35*). Although the function of BMP signal on Bam regulation in male GSCs is less clear (*13*), it also appears to be required for GSC maintenance at least in part, by repressing bam expression (*11*). Therefore, we next wondered whether Dpp signal has the opposite effect on Bam expression in GSCs as compared to differentiating cells.

If the Dpp signal acts on Bam in differentiating cells to inhibit de-differentiation, Dpp should enhance Bam expression. Thus, blocking Dpp diffusion would be expected to show reduction in Bam expression in differentiating cells. To test this, we blocked Dpp diffusion using Morphotrap (*fasIII>nrvMT in mGL-dpp* homozygous background, rescued by *pPA dpp 8391/X*) and stained these testes for Bam protein. As we expected, Bam expression was reduced in 4-to-8 cell SGs and tended to reach a peak in 16-cell SGs, in contrast to the control in which the peak of Bam expression was seen at 8-cell SGs (Figure 6A-E) as reported previously (*36*), suggesting that Mad promotes upregulation of Bam in GB-SGs upon exit from GSC stage, opposite of its inhibitory function in GSCs (Figure 6A-D, arrowheads and F, and Figure S6A-J, arrowheads). Supporting our hypothesis that Dpp signal have dual role on *bam* expression, germline tumors induced by overexpression of the constitutive active form of Tkv (nos>TkvCA) are a mixture of Bam positive and negative cells as reported previously (Figure 6G) (*13*).

**Figure 6.**
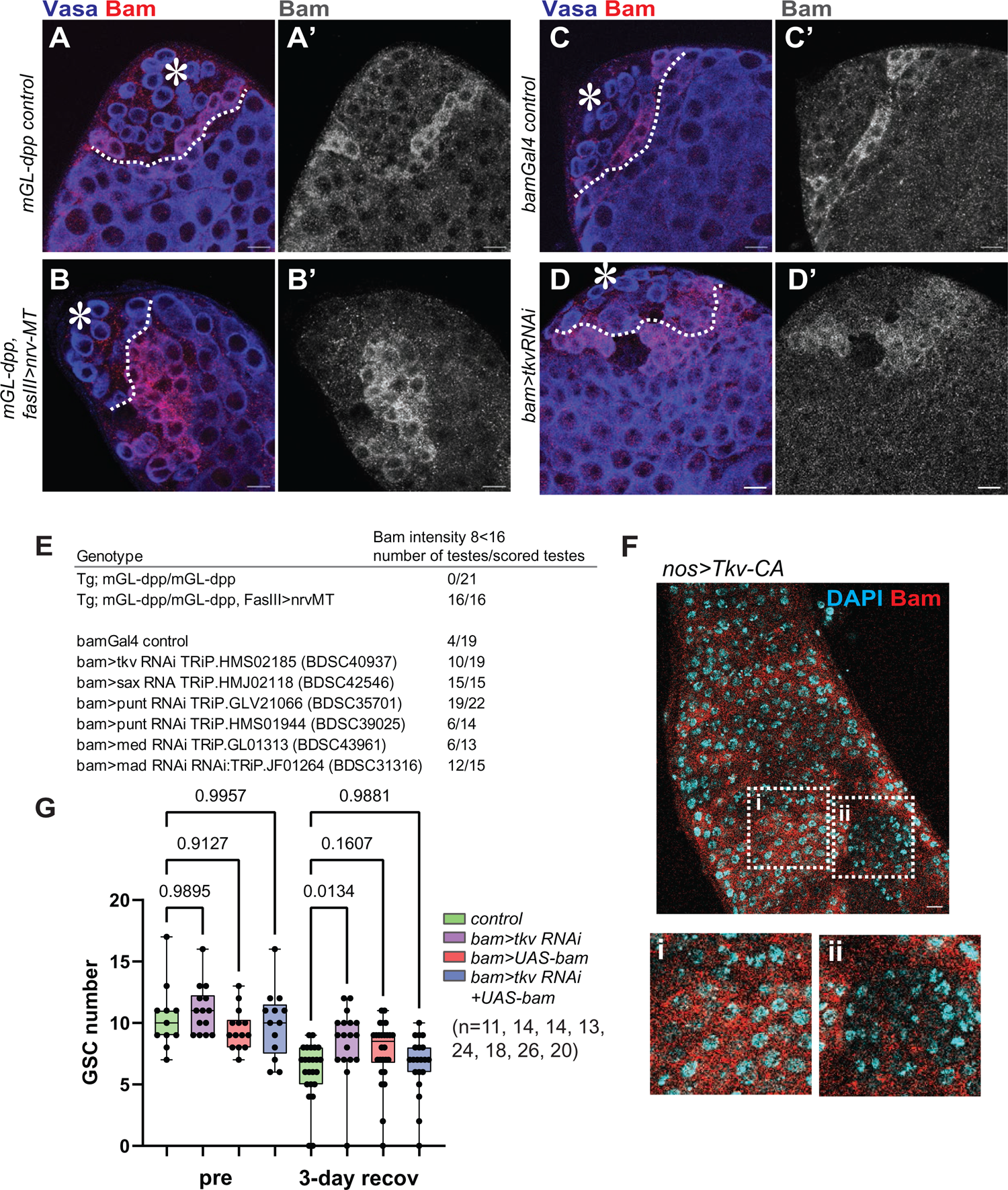
Diffusible fraction of Dpp activates Bam expression in differentiating germ cells to prevent de-differentiation. **A, B**) Representative Bam staining images of *mGL-dpp* testis tip without (**A**) or with (**B**) fasIIIGal4 driven Nrv-MT expression. **C, D**) Representative Bam staining images of testis tip without (**C**) or with (**D**) bamGal4 driven tkvRNAi. In A-D, boundary between 8-cell SGs and 16-cell SGs are divided by white broken lines. **E**) Frequency of testes in which Bam shows higher expression level in 16-cell stages than 8-cell stages (see *Method* for details of scoring) for indicated genotypes. Tg indicates transgene located on X chromosome (*pPA dpp 8391/X*) for rescuing semi-lethal *mGL-dpp* allele. **F**) Representative images of the cells in tumor with Bam staining in the testis of flies expressing Tkv-CA under the nosGal4 driver. Right panels are magnification of squared regions of i and ii in the image. **G**) Changes in GSC number during recovery from forced differentiation of GSCs in indicated genotypes. P-values were calculated by Šídák’s multiple comparisons test. All scale bars represent 10 μm. Asterisks indicate approximate location of the hub. Fixed samples were used for all images. Box plots show 25–75% (box), minimum to maximum (whiskers) with all data points. “n” indicates the number of scored testes in **G**.

To test whether reduced Bam levels is the cause of accelerating de-differentiation, we attempted to rescue the bam>Tkv RNAi phenotype by combining it with Bam overexpression. Strikingly, bamGal4 mediated expression of Bam abrogated the observed enhancement of de-differentiation in bam>Tkv RNAi alone (Figure 6H), indicating that Dpp signal outside of the hub inhibits de-differentiation through augmentation of Bam expression.

### Mad switches its function on the *bam* promoter

So far, our data suggest that Dpp/Gbb-Tkv/Punt or Sax/Punt act on Mad/Med and these downregulate Bam expression in GSCs, whereas it upregulates Bam expression in GB/SGs. How does the same pathway exert the opposing effects on same target gene *bam*?

In the testis, Dpp signal is highest in GSCs, with strong pMad intensity which becomes substantially lower in GB/SGs (Figure 2I). Therefore, we wondered if the opposing output of Dpp depends on pMad concentration. To mimic GSC’s pMad level in differentiating cells, we overexpressed Mad under the bamGal4 driver (bam>Mad). If high concentration of Mad reverse the function of low concentration of Mad, we should observe Mad overexpression has opposite effect on dedifferentiation (i,e, enhancing the dedifferentiation). However, Mad overexpression in hs-Bam flies reduced recovery rate of GSCs after heat-shock mediated GSC depletion (Figure 7A), suggesting that functional switch of Mad from repressor to activator unlikely depends on concentration of pMad, and other factor(s) may regulate Mad function in a stage specific manner.

**Figure 7.**
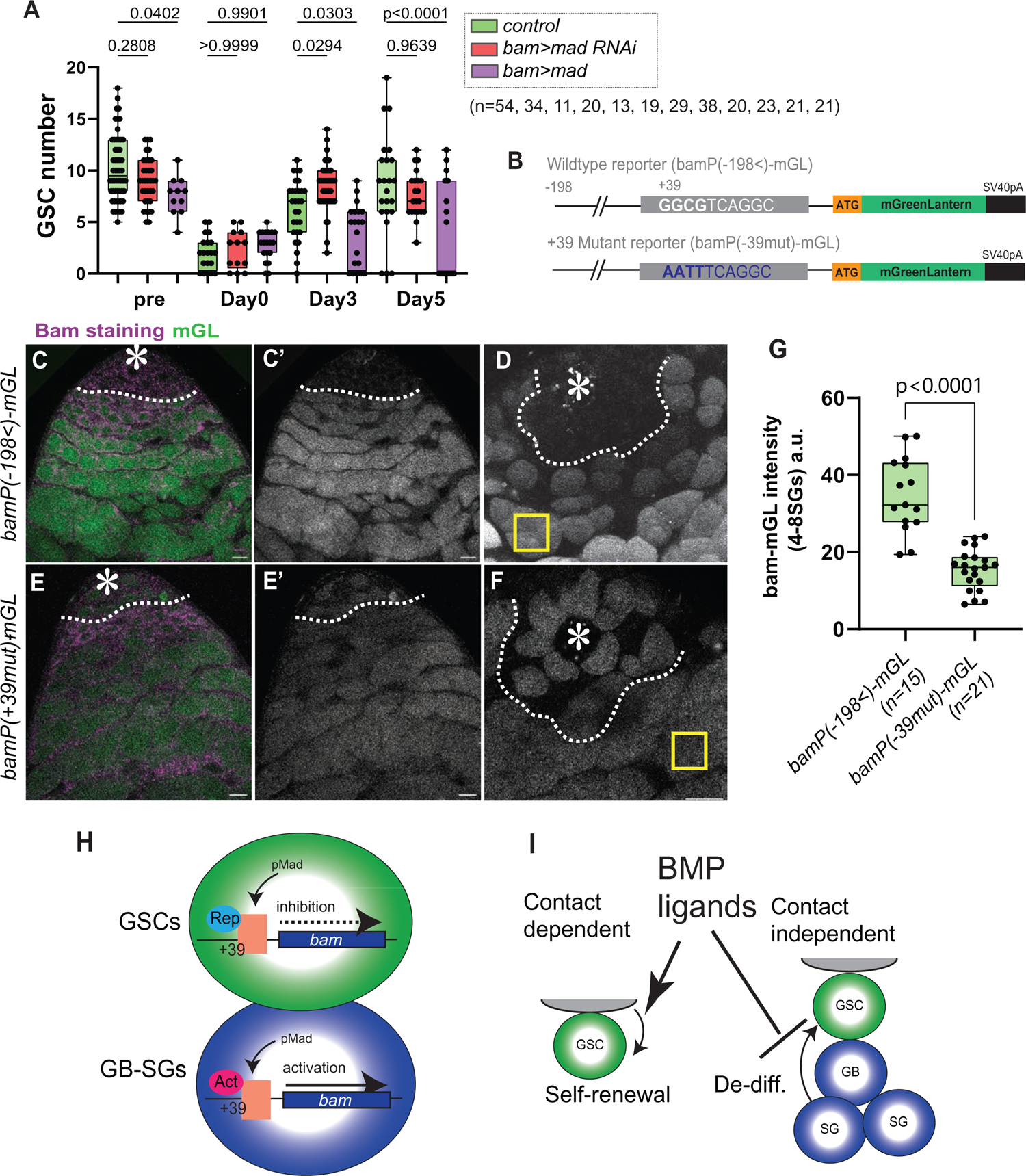
Mad switches its function on the *bam* promoter. **A)** Changes in GSC number during recovery from forced differentiation of GSCs without or with overexpression of Mad (UAS-Mad), knockdown of Mad (Mad RNAi) under the bamGal4 driver. **B)** Structure of *bam* promoter-mGL reporter construct. The previously reported (*37*) core Mad binding element, +39 to +44 GGCGTC is shown in white letters and were mutated for mutant reporter constructs (GGCG to AATT). **C-F)** Representative images in the testis of flies harboring indicated bamP (promoter)-mGL reporters. **D** and **F**) show examples of mGL signal in live tissue used for quantification of intensities in 4-8-cell SGs (yellow squares). White broken lines indicate boundary between 2- and 4-cell SGs where Bam staining turns negative to positive. Wild type reporter, bamP(−198<)-mGL, shows mGL signal only distal from the line (**D**), whereas mutant reporter, bamP(+39mut)-mGL shows mGL signal both apical and distal areas (**F**). +39 mut reporter shows lower mGL intensity in 4-8-cell SGs. **G**) Quantification of reporter intensity in 4-8 cell SGs of indicated reporters. **H**) Model. pMad is required for suppression of bam expression in GSCs, whereas pMad is required for upregulation of bam in GB and SGs both through +39 Mad binding site. Functional switch of pMad may be dependent on co-factors differently associate with pMad in GSCs and GB-SGs (Rep: repressor, Act: activator). **J**) Model. Niche ligands, Dpp and Gbb have effect on GSCs in a contact dependent manner and on differentiating germ cells (GBs and SGs) through diffusion. Contact dependent signal is required for stem cell maintenance (Self-renewal), whereas diffusing ligands promote differentiation of daughter cells via preventing de-differentiation (De-diff). All scale bars represent 10 μm. Asterisks indicate approximate location of the hub. Fixed samples were used for **A, C** and **E**. Live tissues were used for **D, F** and **G**. P-values were calculated by Šídák’s multiple comparisons tests for **A** and student-t tests for **G** and provided on each graph. Box plots show 25–75% (box), minimum to maximum (whiskers) with all data points. “n” indicates the number of scored testes.

Previously, Mad/Med complex has been shown to directly bind to silencer element within bam promoter to suppress its expression (*35*). Therefore, we next wondered whether Mad directly act on bam promoter both in GSCs and GB/SGs to exert opposite roles. If so, the Mad binding domain in the silencing element of *bam* promoter may alter its function from a repressor to an activator. To test this possibility, we generated bam promoter reporter constructs harboring same mutation on +39 site as previously described (Figure 7B) (*37*). We observed expression of +39 site-mutated reporter in GSCs, indicating that this site is required for suppression of bam expression in GSCs similar to female GSCs (*37*). In addition, we noticed that +39 site-mutated-reporter shows drastically lower intensity relative to the control reporter specifically in SGs (Figure 7D-G), indicating that the +39 site is required for upregulation of *bam* in SGs opposed to its silencer effect in GSCs, and that Mad acts oppositely on +39 site of bam promoter in GSCs and in SGs (Figure 7H). Mad has been shown to interact with numerous co-factors to be either a transcriptional repressor or activator (*38*). Further studies will be required to identify co-factors that associate with Mad specifically in a stage specific manner.

Taken together, this study provides a new paradigm of niche space restriction. We provide clear evidence that a soluble niche ligand can spread from a niche and facilitates differential signaling responses in stem cells versus their differentiating progenies (Figure 7J).

## Discussion

In this study, we demonstrate the presence of a diffusible fraction of Dpp and show that it has a key function outside the niche in promoting GSC daughter cells differentiation, a role opposite to its function in the niche in promoting GSC self-renewal. These opposing signaling outcomes are achieved by the same canonical BMP pathway, via the receptor Tkv, Sax and Punt and downstream effector Mad/Medea, which represses Bam expression in stem cells but upregulates Bam expression in differentiating cells. It has been suggested that the Dpp has only a minor effect in GSC maintenance in the testicular niche based on the mild stem-cell loss phenotype of *dpp* mutants (*11*). This may be accounted by opposed function of Dpp in GSC and GB/SGs. Because Dpp mutants can cause GSC loss and enhanced de-differentiation at the same time, it results in apparently normal GSC number remained in the niche.

The niche Dpp has been postulated to act as a highly localized signal as pMad can be observed exclusively in GSCs and their only immediate progenies. However, our work suggests that the distribution of Dpp is rather uniform, implying that the signal reception is not uniform. Indeed, many works have revealed how a steep gradient of BMP response is established within just one cell diameter (*39–50*). Many of these studies postulated redundant mechanisms in which either stem cells enhance the signal reception, or differentiating cells actively suppress it.

Alternatively, specific composition of ligands may not be so uniform. Our study suggests a requirement of both Dpp and Gbb for observed cellular responses, and co-localization of Dpp-Gbb was exclusively seen in the hub (Figure 5L, M), suggesting a possibility that Dpp-Gbb may form heterodimer and distribution of which might be tightly restricted around the hub even though each ligand alone can diffuse freely. A recent report has demonstrated that the heterodimers of Dpp-Gbb preferentially produced in Dpp producing cells and play critical roles during development (*31, 51*). This may contribute to the formation of sharply graded signaling outcomes around the niche. It would be interesting to investigate the precise distribution and composition of these ligands in a quantitative manner.

Because mammalian homologs of Dpp, the TGF-beta family genes, broadly regulate processes in many types of stem cell niches (*9*), we propose that the diffusion of the ligands may be a common mechanism in stem cell niches to ensure their spatial restriction and asymmetric outcome of stem cell division. Intriguingly, differential effects for a BMP ligand, transforming growth factor (TGF)-β, albeit focused on proliferation, have been reported in hematopoietic stem cell (HSC) niche, where low concentrations of TGF-β induces proliferation of myeloid-biased HSCs but inhibit proliferation of lymphoid-biased HSCs (Ly-HSCs) (*52, 53*). In this case, it is still unknown whether the ligand forms a gradient around HSC niche and whether these progenitors are located distinct positions that subject them to different TGF-β fractions. The elucidation of the basis of these differential outputs based on ligand behavior is a fascinating future study.

## Acknowledgements

We thank Markus Affolter, Margaret T. Fuller for discussion and suggestions. Yukiko Yamashita, Michael Buszczak, Thomas Kornberg, Ryo Hattori, and the Bloomington Drosophila Stock Center and the Developmental Studies Hybridoma Bank for reagents; Marie Bao (Life Science Editors) for manuscript editing. This research is supported by R35GM128678 from the National Institute for General Medical Sciences and a start-up fund from UConn Health (to M.I.) and an SNSF Ambizione grant (PZ00P3_180019) (to S.M.).

## Author Contributions

M.I. conceived the project. M.I., S.M.R., A.T. and E.K.B.,designed and executed experiments and analyzed data. S.M. generated *mGL-dpp* line provided all morphotrap and tagged dpp or gbb lines, assisted with the design of the experiments. S.P. generated bam reporter flies. M.B.B. generated *UASp-bam* transgenic fly. A.E.C., quantitative analysis and interpretation of imaging data. M.I. and M.A. drafted manuscript. All authors edited the manuscript.

## Data availability statement

The data that support all experimental findings of this study are available within the paper and its Supplementary Information files. All imaging files will be available from the corresponding author M.I. upon request.

## Software and code

Confocal images were collected by a Zeiss LSM 800 Confocal microscope using ZEN software. Confocal images were analyzed using ImageJ/Fiji software (version 2.1.0). Statistical analysis and figures were generated using GraphPad Prism software (version 9.2.0). Raw data necessary to reproduce all statistical analyses and results in the paper are provided in the source data file provided with this paper.

## Materials and Methods

### Fly husbandry and strains

Flies were raised on standard Bloomington medium (Lab express) at 25°C (unless temperature control was required). The following fly stocks were obtained from Bloomington stock center (BDSC); *nosGal4* (BDSC64277); *hs-bam* (BDSC24636); *tkv RNAi* (BDSC40937); Nrv1 morphotrap (*lexAop-UAS-GrabFP.B.Ext.TagBFP*, BDSC68173); mCD8-morphotrap *(lexAop-UAS-morphotrap.ext.mCh*, BDSC68170); *medea RNAi:TRiP.GL01313* (BDSC43961); *mad RNAi:TRiP.JF01264* (BDSC31316); *sax RNAi:TRiP.HMJ02118* (BDSC42546); *punt RNAi: TRiP.HMS01944* (BDSC39025); *punt RNAi: TRiP.GLV21066* (BDSC35701); *gbb RNAi:TRiP.HMS01243* (BDSC34898); *tkv-CA* (BDSC36537); *UAS-GFP.dsRNA.R* (BDSC44415); *gbb-GFP.R* (BDSC63055). *yw* (BDSC189) was used for wildtype. *UAS-GFP-Mad* (*50*)*, HA-dpp* (*30*)*, UAS-HA-trap* (*30*)*, HA-gbb* (*31*) *dppGal4* (*30*) lines are described elsewhere. *FasIIIGal4* was obtained from DGRC, Kyoto Stock Center (A04-1-1 DGRC#103-948). *GFP-dpp* and *mCherry-dpp* (FBst0086273) knock-in lines was kind gift from Thomas Kornberg and Ryo Hattori (*54*). *pVas-Vasa-mCherry* (FBtp0065762) (*55*), *UAS-histone H3-GFP* and *bamGal4* on 3^rd^ was kind gifts from Yukiko Yamashita.

Temperature shift was performed by culturing flies at room temperature and shifted to 29°C upon eclosion for the 4 days before analysis. Combinations of Tub-Gal80ts (a gift from Cheng-Yu-Lee) with c587Gal4 (a gift from Yukiko M. Yamashita) were used.

For all crosses for obtaining *mGL-dpp* homozygous flies, transgenic allele containing *dpp* locus (*pPA dpp 8391/X*) (*21*) was introduced to assist embryonic expression and rescue semi-lethality. This transgene has been known to only rescue early development of *dpp* null mutant (*21*).

### Generation of *mGL-dpp* and *mSC-dpp* alleles

The detail procedure to generate endogenously tagged *dpp* alleles were previously reported (*30*). In brief, utilizing the *attP* sites in a MiMIC transposon inserted in the *dpp locus* (*MiMIC dppMI03752*, BDSC36399), about 4.4 kb of the *dpp* genomic sequences containing the second (last) coding exon of *dpp* including a tag and its flanking sequences was inserted in the intron between *dpp*’s two coding exons. The endogenous exon was then removed using FLP-FRT to keep only the tagged exon. mGL (mGreenLantern (*56*)) or mSC (mScarlet (*57*)) were inserted in frame after amino acid 485 (NM_164488.2) after the last processing site to tag all the Dpp mature ligands. mGL coding sequences after the last processing site. The detail characterization of these alleles are described in (*22*).

### Generation of *UASp-bam* transgenic line

*bam* cDNA was PCR-amplified from cDNA pool isolated from wild-type testis (*yw*) using the following primers with restriction sites (underlined): NotI bam Forward 5’-AC<u>GCGGCCGC</u>ACCATGCTTAATGCACGTGACGTGTGTC-3’ AscI bam Reverse 5’-AT<u>GGCGCGCC</u>TTAGCTTCTGAAGCGAGGTACACGTCCGG-3′PCR products were then digested with NotI and AscI and ligated to a modified pPGW vector (kind gift from Michael Buszczak) using NotI and AscI sites within the multiple cloning site and verified by Sanger sequencing (Genewiz). Transgenic flies were generated using strain attP2 by PhiC31 integrase-mediated transgenesis (BestGene).

### Generation of *bam* reporter transgenic lines

Bam promoter fragment was amplified from genomic DNA using following primers.Overlap sequences for Gibson Assembly reaction were added for each primer.

1. Bam promoter-F: 5’AGCGGATCCAAGCTTGCATGCGGTACCCCAAATCAGTGTGTATAATT-3′
2. Bam promoter-R: 5’-TATTCTTAAGTTAAATCACACAAATCACTCGAT-3′

Mad binding site mutation was designed as previously described (*35*) and introduced by PCR using following primers. mut-F: 5’-CGCAGACAGCGTAATTTCAGCGATTTCAAACGGTAAAAAG-3′ mut-R: 5’-GAAATTACGCTGTCTGCGAATTCAGGAGAAAGAGGAAGAA-3′Bam promoter-F/Bam promoter-R fragment was assembled with pUAST-GFP-attB vector (gift from Cheng-Yu-Lee) digested by SphI/NotI to remove UAS promoter located between these sites and mGL fragment.

For Mad-binding site mutant reporter, Bam promoter-F/mut-R, mut-F/Bam promoter-R fragments were assembled with the same vector and mGL fragment. mGL fragment was amplified from synthesized DNA (below) by using following primers mGL-forward: 5’-AACTTAAGAATAATGGTGAGCAAGGGCGAGGAGCTGT-3’ mGL-reverse: 5’-TAGAGGTACCCTCGAGCCGCTTACTTGTACAGCTCGTCCATGCCGAGA-3’ mGL gBlock fragment: 5’atggtgagcaagggcgaggagctgttcaccggggtggtgcccatcctggtcgagctggacggcgacgtaaacggccacaagttcagcgtccgcggcgagggcgagggcgatgccaccaacggcaagctgaccctgaagttcatctgcaccaccggcaagctgcccgtgccctggcccaccctcgtgaccaccttaggctacggcgtggcctgcttcgcccgctaccccgaccacatgaagcagcacgacttcttcaagtccgccatgcccgaaggctacgtccaggagcgcaccatctctttcaaggacgacggtacctacaagacccgcgccgaggtgaagttcgagggcgacaccctggtgaaccgcatcgtgctgaagggcatcgacttcaaggaggacggcaacatcctggggcacaagctggagtacaacttcaacagccacaaggtctatatcacggccgacaagcagaagaacggcatcaaggctaacttcaagacccgccacaacgttgaggacggcggcgtgcagctcgccgaccactaccagcagaacacccccatcggcgacggccccgtgctgctgcccgacaaccactacctgagccatcagtccaaactgagcaaagaccccaacgagaagcgcgatcacatggtcctgaaggagagggtgaccgccgccgggattacacatgacatggacgagctgtacaagtaa3’

The amplified fragments were assembled using Gibson Assembly kit (NEB) and verified by Sanger sequencing (Genewiz). Transgenic flies were generated using strain attP40 by PhiC31 integrase-mediated transgenesis (BestGene).

All gBlock fragments and primers used in this study were synthesized by Integrated DNA Technologies (IDT).

### Induction of de-differentiation

Induction of de-differentiation was performed following previously described method with modifications (*25*). Approximately 0- to 3-day-old adult flies carrying hs-Bam (BDSC24636) transgene were raised in 22°C and heat-shocked in a 37°C water bath for 30 min twice daily in vials with fly food. Vials were placed in a 29°C incubator between heat-shock treatments. After 6-time treatments, vials were returned to 22°C for recovery. Testes were dissected at desired recovery time points.

### Short-term live imaging

Testes from newly eclosed flies were dissected into Schneider’s Drosophila medium containing 10% fetal bovine serum and glutamine–penicillin–streptomycin. These testes were placed onto Gold Seal Rite-On Micro Slides’ 2 etched rings with media, then covered with coverslips. Images were taken using a Zeiss LSM800 airyscan with a 63× oil immersion objective (NA = 1.4), with 10-20 z-stacks (interval 1µm). within 30 minutes. For short-term live imaging experiments, imaging was performed within 30 minutes.

### Long-term live imaging of de-differentiation

Imaging was performed as previously described (*58*). The testes were dissected in 1X Becker Ringer’s solution (*58*) and then mounted onto a 35mm Glass Bottom Dishes (Nunc). 500 μL of 1 mg/mL poly-L-lysine (Sigma) was pipetted onto the coverslip portion of the imaging dish and incubated for 5- to 7-hours at room temperature. Then, poly-L-lysine solution was replaced to the Becker Ringer’s solution and testes were mounted onto poly-L-lysine layer with the tip of the testes oriented toward bottom. Next, Becker Ringer’s solution was slowly removed and replaced with 3ml of room temperature Schneider’s *Drosophila* medium supplied with 10% fetal bovine serum and glutamine–penicillin–streptomycin (Sigma). Z-stacks (2µm interval, for 11 stacks) were taken using Zeiss LSM800 airyscan, 1AU-pinhole with 63X oil immersion objective (NA = 1.4) every 10 minutes for overnight (16 hours). Preset tiling function (Zen software, Zeiss) was used for sequential imaging of multiple positions to obtain time-lapse images from 5-to-8 testes per night.

### Immunofluorescence Staining

Testes were dissected in phosphate-buffered saline (PBS) and fixed in 4% formaldehyde in PBS for 30–60 minutes. Next, testes were washed in PBST (PBS + 0.2% TritonX-100, Thermo Fisher) for at least 60 minutes, followed by incubation with primary antibody in 3% (or 5% for pMad staining) bovine serum albumin (BSA) in PBST at 4°C overnight. Samples were washed for 60 minutes (three times for 20 minutes each) in PBST, incubated with secondary antibody in 3% BSA in PBST at room temperature for 2 hours and then washed for 60 minutes (three times for 20 minutes each) in PBST. Samples were then mounted using VECTASHIELD with 4’,6-diamidino-2-phenylindole (DAPI) (Vector Lab). For pMad staining, testes were incubated with 5% BSA in PBST for 30min at room temperature prior to primary antibody incubation to reduce background.

The primary antibodies used were as follows: rat anti-Vasa (RRID: AB_760351, 1:20; DSHB); mouse anti-Hts (1B1; RRID: AB_528070, 1:20; DSHB); mouse-anti-FasIII (RRID:AB_528238, 1:20, 7G10; DSHB); mouse anti-γ-Tubulin (GTU-88; RRID:AB_532292, 1:400; Sigma-Aldrich); Rabbit anti-pMad (RRID:AB_491015, 1:300; Cell Signaling Technology, Cat# 9516); Mouse anti-phospho-Histone H3 (Ser10), clone 3H10 (RRID:AB_477061; 1:200, Sigma-Aldrich); Rabbit anti-HA C29F4 (RRID:AB_1549585, 1:300, Cell Signaling Technology, Cat# 3724). AlexaFluor-conjugated secondary antibodies (Abcam) were used at a dilution of 1:400.

For Bam staining, 0.2% Tween-20 (Thermo Fisher) was used instead of TritonX-100 for PBS-T. mouse anti-Bam (1:20) antibody was a kind gift from Michael Buszczak. Images were taken using Zeiss LSM800 confocal microscope with airyscan module by using 1AU-pinhole with 63X oil immersion objective (NA = 1.4). Images were processed by image J/FIJI.

### Chloroquine treatment

Testes from newly eclosed flies were dissected into Schneider’s Drosophila medium containing 10% fetal bovine serum and glutamine–penicillin–streptomycin with or without 100 μM chloroquine (Sigma) and incubated for 4 hours at room temperature. These testes were placed onto Gold Seal Rite-On Micro Slides’ 2 etched rings with media, then covered with coverslips. An inverted Zeiss LSM800 airyscan with a 63× oil immersion objective (NA = 1.4) was used for imaging.

### FRAP analysis

Testes from newly eclosed flies were dissected into Schneider’s *Drosophila* medium containing 10% fetal bovine serum and glutamine–penicillin–streptomycin. These testes were placed onto Gold Seal Rite-On Micro Slides’ 2 etched rings with media, then covered with coverslips. Images were taken using a Zeiss LSM800 confocal microscope with a 63× oil immersion objective (NA = 1.4) within 30 minutes. For all live imaging experiments, imaging was performed within 30 minutes. Fluorescence recovery after photo-bleaching (FRAP) of mGL Dpp signal was undertaken using a Zeiss LSM800 with airyscan module by using 1AU-pinhole with 63X oil immersion objective (NA = 1.4). Zen software was used for programming each experiment. Encircled areas of interest (randomly chosen 5µm-diameter circles from the area within less than 40 µm away from the testis tip) were photobleached using the 488 nm laser (laser power; 100%, iterations; 10). Fluorescence recovery was monitored every 10 seconds for single z-plane. Background signal taken in outside of the tissue in each time point were subtracted from the signal of bleached region. Dextran dye permeabilization assay was performed as described previously (*59*). Briefly, testes were incubated with 10kDa dextran conjugated to AlexaFluor 647 (Thermo Fisher, Catalog number: D22914) at a final concentration of 0.2μg/μl in 1 mL media for 5min then immediately subjected for imaging within 30min. Acquisition setting was adjusted for each sample and normalized by calculating % recovery rate.

Images were processed by image J/FIJI. %recovery rate was calculated as follows; Let I^t^ be the intensity at each time point (t), I^post^ be the intensity at post-bleaching (first postbleach scan) and I^pre^ be the intensity at pre-bleaching. The governing equation of % recovery is: % recovery= (I^t^ – I^post^)/(I^pre^ - I^post^) x100.

### Quantification of pMad intensities

Image-J/Fiji software was used for image quantification. Average intensity was measured for anti-pMad staining from each GSC nucleus using a single slice taken by 1AU-pinhole with 63X/1.4 NA oil objective confocal imaging, and background level measured distal region of the same testis was subtracted. Same acquisition setting was used across the samples. To normalize the staining conditions, the average intensities of pMad from four cyst cells (CCs) in the same testes were used as internal control and the ratios of intensities were calculated as each GSC per average intensities of CC. The means and s.d. were plotted to the graph for each genotype. Mean intensity values (a.u.) of CCs were unchanged for genotypes shown in Figure S2D-G (see details in main text).

### Quantification and staging of Bam expression and Bam reporter intensities

Image-J/Fiji software was used for quantification. Average intensity was measured for anti-Bam staining or mGL signal from regions of 4- or 8-cell cysts or 16-cell cysts using a single z-stack (1µm interval) taken by 1AU-pinhole with 63X/1.4 NA oil objective airyscan imaging, and subtracted background measured from distal area of the testis within the same slice. Same acquisition setting was used across the samples. For Figure 6E, average intensities of measurement of 3 portions each from 8-cell cysts and 16-cell cysts were measured and subtracted the background (taken from distal area of the same testis), then plotted to calculate the ratio of 16SG intensity/8SG intensity for each testis. When the ratio was greater than 1, the testis was categorized for “Bam intensity 8<16”.

### Scoring of centrosome and spindle orientation

Centrosome misorientation was indicated when neither of the two centrosomes were closely associated with the hub-GSC interface during interphase. Spindle misorientation was indicated when neither of the two spindle poles were closely associated with the hub-GSC interface during mitosis.

### Statistical analysis and graphing

No statistical methods were used to predetermine sample size. The experiments were not randomized. The investigators were not blinded to allocation during experiments and outcome assessment. All experiments were independently repeated at least 3 times to confirm the results. Statistical analysis and graphing were performed using GraphPad Prism 9 software.

### Supplemental Data

Individual numerical values displayed in all graphs are provided.

