## supplementary materials for "Diffusible fraction of niche BMP ligand safeguards stem-cell differentiation"

**The PDF file includes:**

Table S1, Figures S1 to S6
Legends for Video S1, Video S2, Source Data

| Alleles | *mCherry-dpp**1 | *GFP-dpp**1 | *mGL-dpp**1 | *mSC-dpp**1 | *HA-dpp**1 | *GFP-dpp* |
|---|---|---|---|---|---|---|
| Sources | Fereres et al., 2019 | Gift from Thomas Kornberg | This study, Rasouliha et al., 2023 | This study, Rasouliha et al., 2023 | Matsuda et al., 2021 | Matsuda et al, 2021 |
| Phenotypes | Homozygous viable<br><br>Potential generation of non-tagged Dpp fraction*2 | Homozygous viable<br><br>Potential generation of non-tagged Dpp fraction*2 | Homozygous semi-lethal<br><br>Rescuable with *pPA dpp 8391/X* | Homozygous semi-lethal<br><br>Rescuable with *pPA dpp 8391/X* | None | Haploinsufficient<br><br>Partially rescuable with *pPA dpp 8391/X* (patterning defects) |
| Tag location | AA465 | AA465 | AA485 | AA485 | AA485 | AA485 |
| GSC phenotypes after expression of MT under hub driver (FasIIIGal4) | Not tested | Reduced pMad with mCD8-MT | Reduced pMad with mCD8-MT | Not tested | Reduced pMad with HAtrap | Not tested |
| De-differentiation phenotypes after expression of MT under hub driver (FasIIIGal4) | Not tested | Accelerated dedifferentiation with Nrv-MT trap | Accelerated dedifferentiation with Nrv-MT trap | Not tested | Not tested | Not tested |

*1 Alleles used in this study

*2 Tags placed at AA465 may be cut out at the last furin processing site

**Table S1. Comparison of fluorescent tagged *dpp* alleles.**

Fluorescent tags are inserted after the indicated amino acid (AA) in *dpp* isoformE (NM_164488.2). Fereres et al., 2019 (53); Rasouliha et al., 2023 (57); Matsuda et al., 2021 (20).

mSC: mScarlet, mGL:mGreen Lantern.

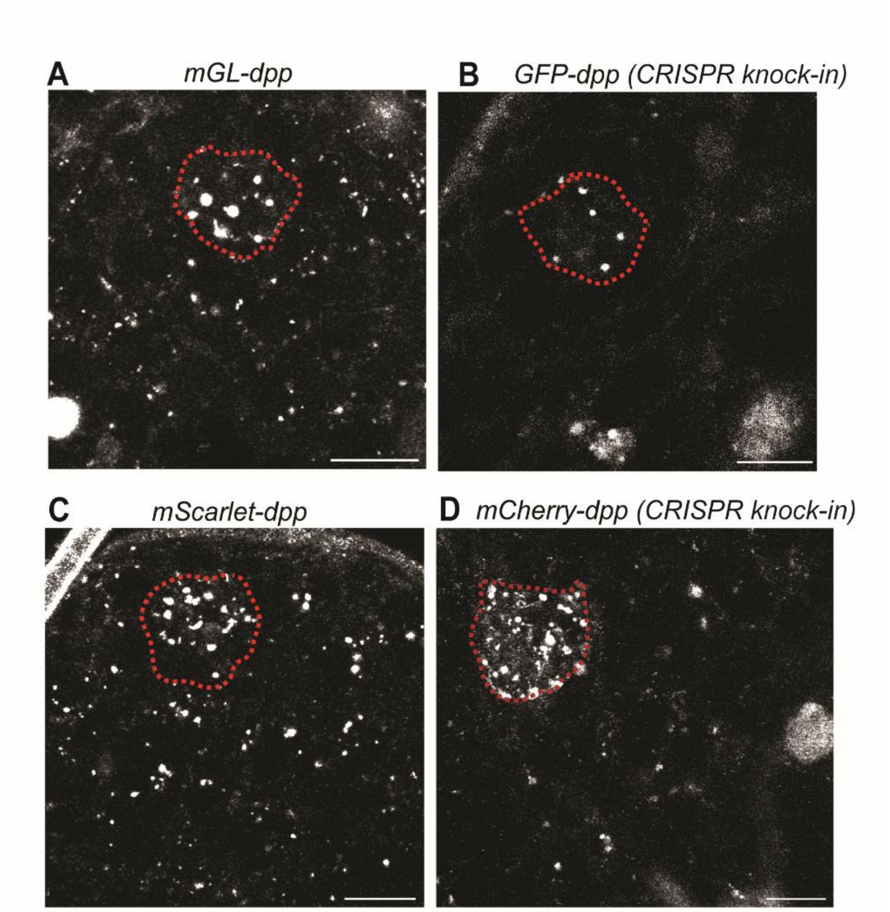


**Figure S1. Comparison of fluorescent patterns of tagged *dpp* alleles.**

Representative images comparing testis tips isolated from homozygous *mGL-dpp* (**A**), homozygous *GFP-dpp* knock-in (**B**), homozygous *mScarlet(mSC)-dpp* (**C**), and homozygous *mCherry-dpp* knock-in (**D**). *mGL-dpp* and *mSC-dpp* homozygous flies were rescued by Tg (*pPA dpp 8391/X*) to assist embryonic development. Scale bars represent 10 μm. The hub is encircled by red broken lines. Live tissues were used for imaging.

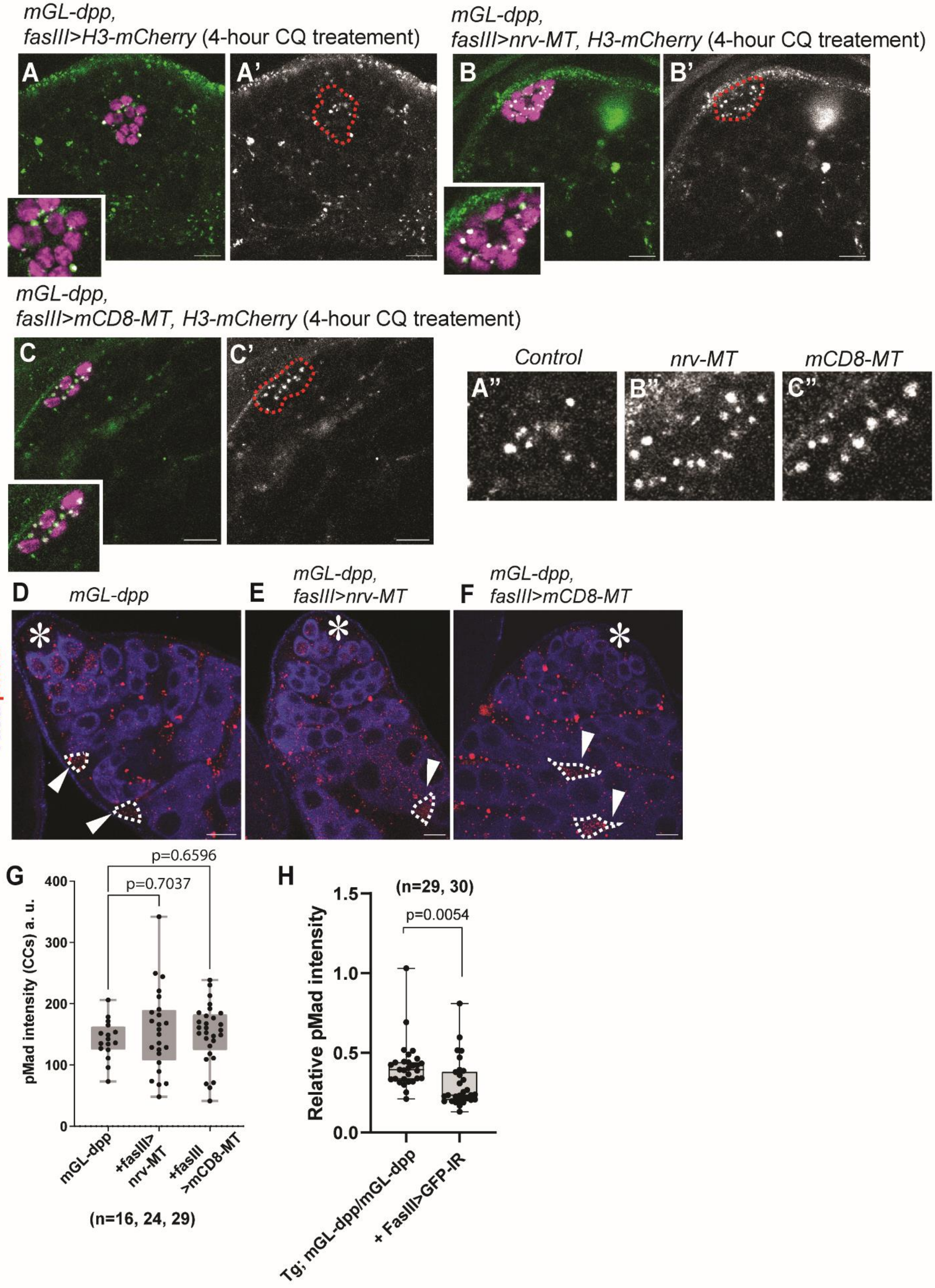
mGL-dpp,
fasIII>H3-mCherry (4-hour CQ treatement)
mGL-dpp,
fasIII>nrv-MT, H3-mCherry (4-hour CQ treatement)
A
A'
B
B'
mGL-dpp,
fasIII>mCD8-MT, H3-mCherry (4-hour CQ treatement)
C
C'
Control
nrv-MT
mCD8-MT
A''
B''
C''
D
mGL-dpp
E
mGL-dpp,
fasIII>nrv-MT
F
mGL-dpp,
fasIII>mCD8-MT
Vasa pMad
G
p=0.6596
p=0.7037
pMad intensity (CCs) a. u.
400
300
200
100
0
mGL-dpp
+fasIII>
nrv-MT
+fasIII
>mCD8-MT
(n=16, 24, 29)
H
(n=29, 30)
p=0.0054
Relative pMad intensity
1.5
1.0
0.5
0.0
Tg; mGL-dpp/mGL-dpp
+ FasIII>GFP-IR

**Figure S2. Characterization of morphotrap phenotypes.**

**A-C**) Representative images of *mGL-dpp* signal after 4-hour CQ treatment in the testes of indicated genotypes. Magnified hub areas are shown in left corner of **A-C**. The hub is encircled by red broken lines in **A'-C'**. Magnified images of hub area are shown in **A"-C"**. Live samples were used. **D-F**) Representative images of pMad staining in CC (marked by broken lines and arrowheads) of indicated genotypes. **G**) Quantification of pMad intensity in CCs of indicated genotypes. P-values were calculated by Dunnett's multiple comparisons tests and provided on the graph. **H**) Quantification of pMad intensity in GSCs (relative to CCs) of fasIIIGal4 driven GFP (mGL) knock-down in *mGL-dpp* homozygous background rescued by Tg (*pPA dpp 8391/X*). pMad signal was significantly reduced in GSCs of GFP RNAi expressing testes, indicating that *pPA dpp 8391/X* is not functional in the hub. The p-values were calculated by student-t-test and provided on the graph. Fixed samples were used for **D-H**.

Asterisks indicate approximate location of the hub. All scale bars represent 10 μm. "n" indicates the number of scored GSCs. Box plots show 25–75% (box), minimum to maximum (whiskers) with all data points.

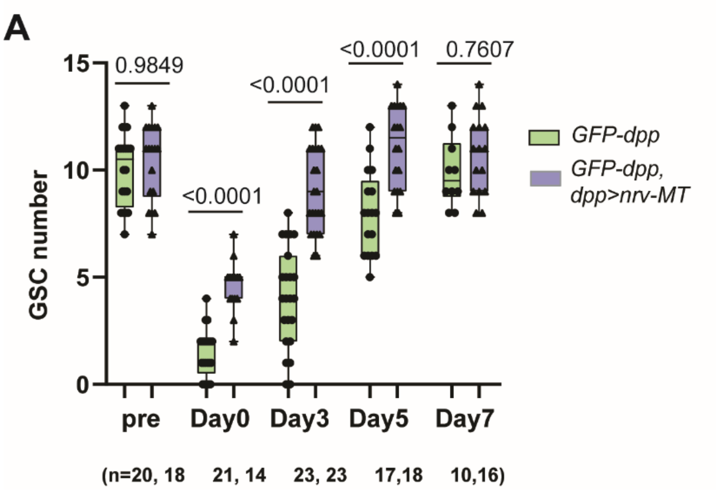


**Figure S3. Morphotrap using alternative genotypes shows similar effect on de-differentiation.**

**A**) Changes in GSC number during recovery from forced differentiation of GSCs. For trapping Dpp, Nrv-MT was expressed under the control of the dppGal4 driver in *GFP-dpp* knock-in homozygous background. *GFP-dpp* homozygous knock-in flies are viable and fertile and were used for the control. P-values were calculated by Šídák's multiple comparisons tests and provided on the graph. Fixed samples were used for scoring. “n” indicates the number of scored testes. Box plots show 25–75% (box), minimum to maximum (whiskers) with all data points.

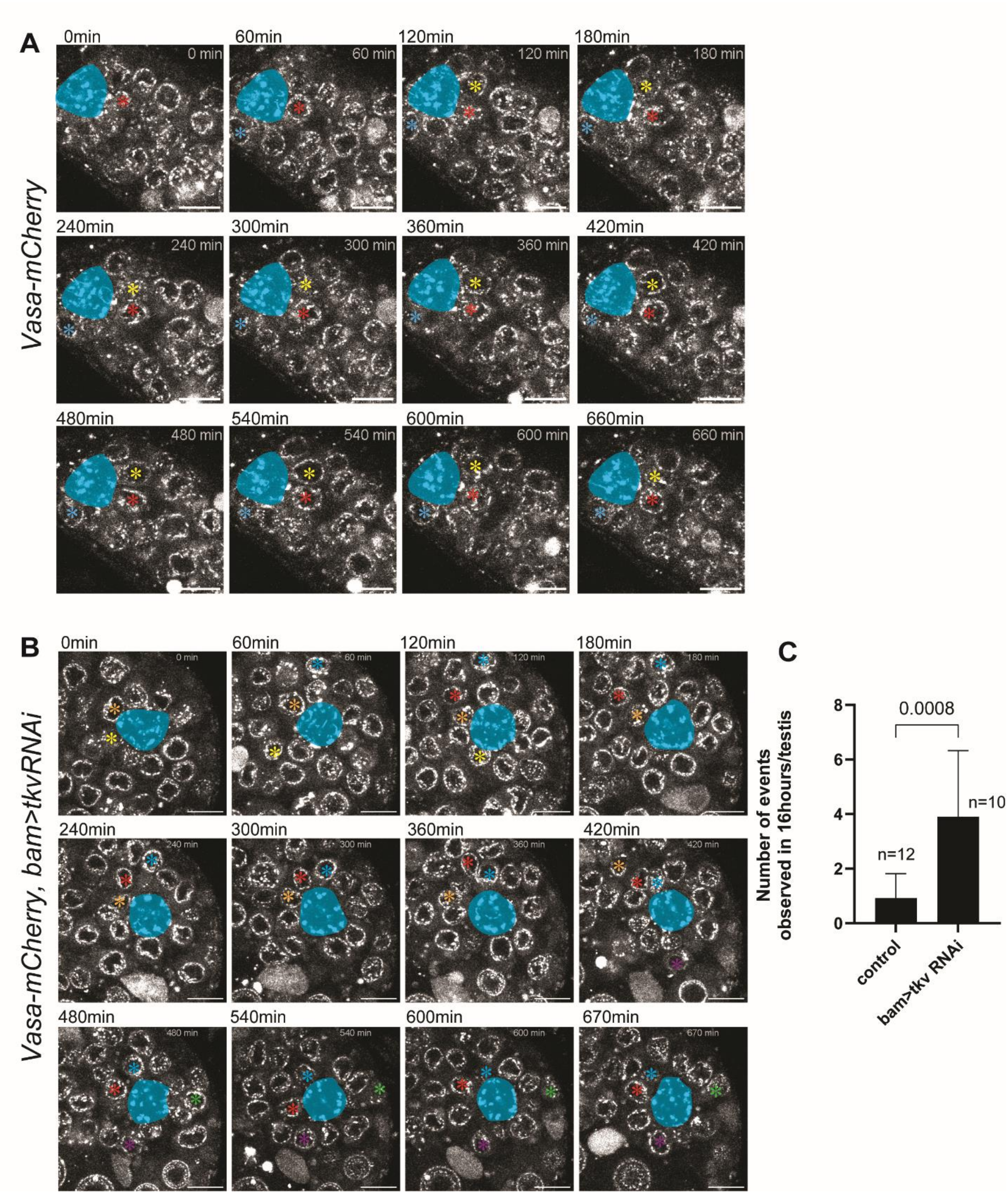
A
0min
60min
120min
180min
240min
300min
360min
420min
480min
540min
600min
660min
Vasa-mCherry
B
0min
60min
120min
180min
240min
300min
360min
420min
480min
540min
600min
670min
Vasa-mCherry, bam>tkvRNAi
C
Number of events observed in 16hours/testis
0.0008
n=12
n=10
control
bam>tkv RNAi

**Figure S4. Live imaging of Tkv knock-down testis shows high frequency of non-asymmetric events.**

**A, B**) Representative time-lapse live series of testis tip of indicated genotypes. Cells traced for entire imaging period are indicated by asterisks in each different color. In A, all marked GSCs stayed attached in the niche throughout the imaging period. In B, all marked cells show non-asymmetric behavior (yellow and orange cells leave from the hub, while other marked cells de-differentiate). Hub area is filled in turquoise blue. All scale bars represent 10 μm. Corresponding movies are provided as supplemental materials (video S1 and video S2). **C**) Number of non-asymmetric events observed in indicated genotypes in 16-hour imaging periods. Vasa-mCherry flies were used for the control. "n" indicates the number of time-lapse series analyzed. Data are means and standard deviation. The p-value was calculated by student-t-test.

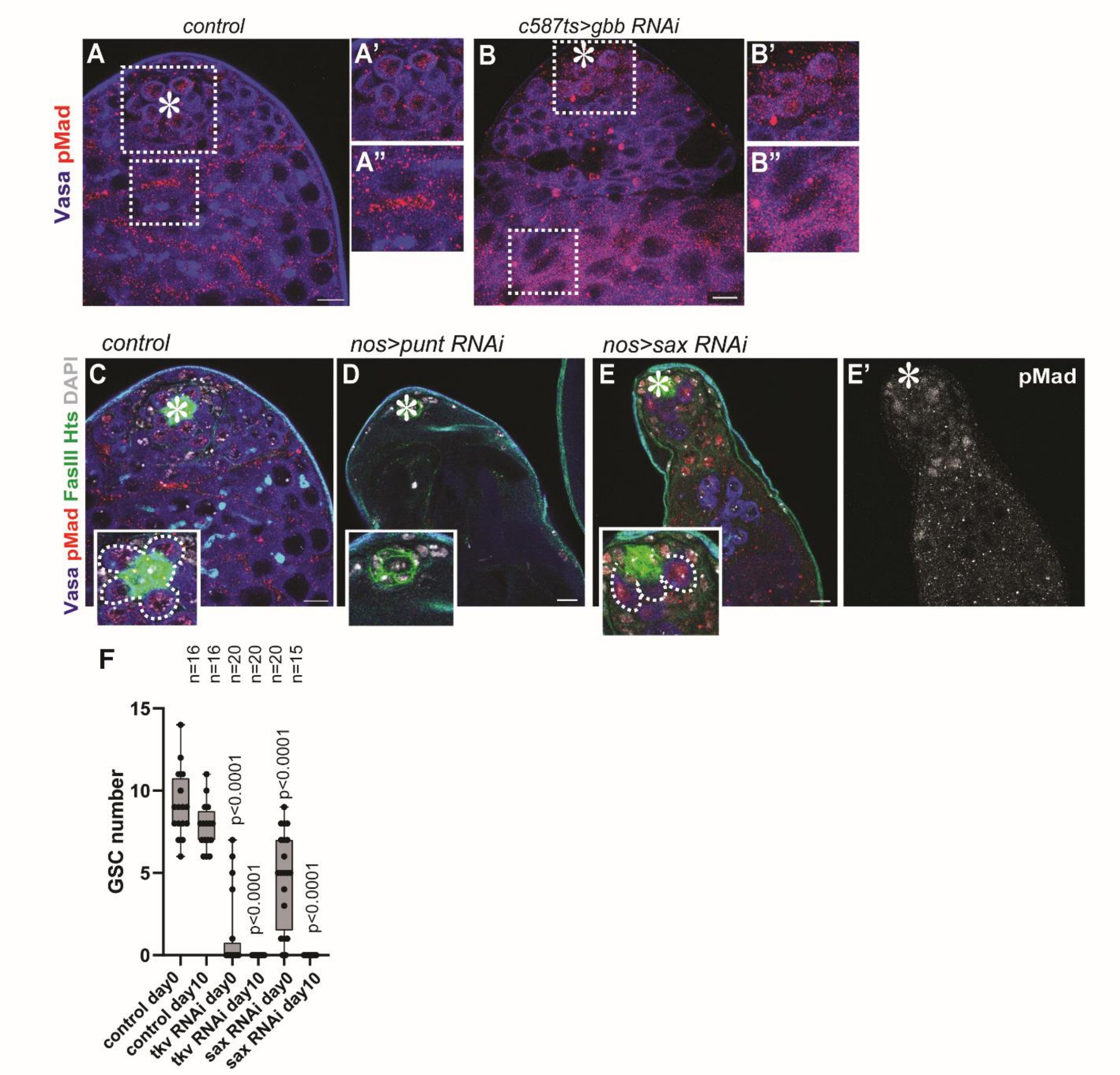


**Figure S5. Sax and Punt show distinct effects on GSC maintenance.**

**A-B)** Representative pMad staining images of testis tips with knock-down of *gbb* under the somatic cell specific driver, c587Gal4$^{ts}$ after 4-days of temperature shift (29 degree). Squared regions in **A, B** are magnified in right panels. **A'**and **B'** show pMad signal in GSCs, which was intact in *gbb* RNAi testes. **A"**and **B"** show pMad signal in CCs, which was not detectable in *gbb* RNAi testes. **C-E)** Representative images of testis tips of no-Gal4 control (**C**) and with knock-down of *punt TRiP.GLV21066* (**D**) or sax *TRiP.HMJ02118* (**E**) under the control of nosGal4 driver. Insets show magnified region around the hub. Punt RNAi shows the testis without any

Vasa positive germ cells. White broken lines encircle GSCs in inset **C** and **E,** showing GSCs with sax RNAi are pMad positive (**E**). **F**) GSC number in indicated age of testes of indicated genotypes. P-values were calculated by Šídák's multiple comparisons tests and provided on the graph. “n” indicates number of scored testes.

Scale bars represent 10 μm. Asterisks indicate approximate location of the hub. Fixed samples were used for all images.

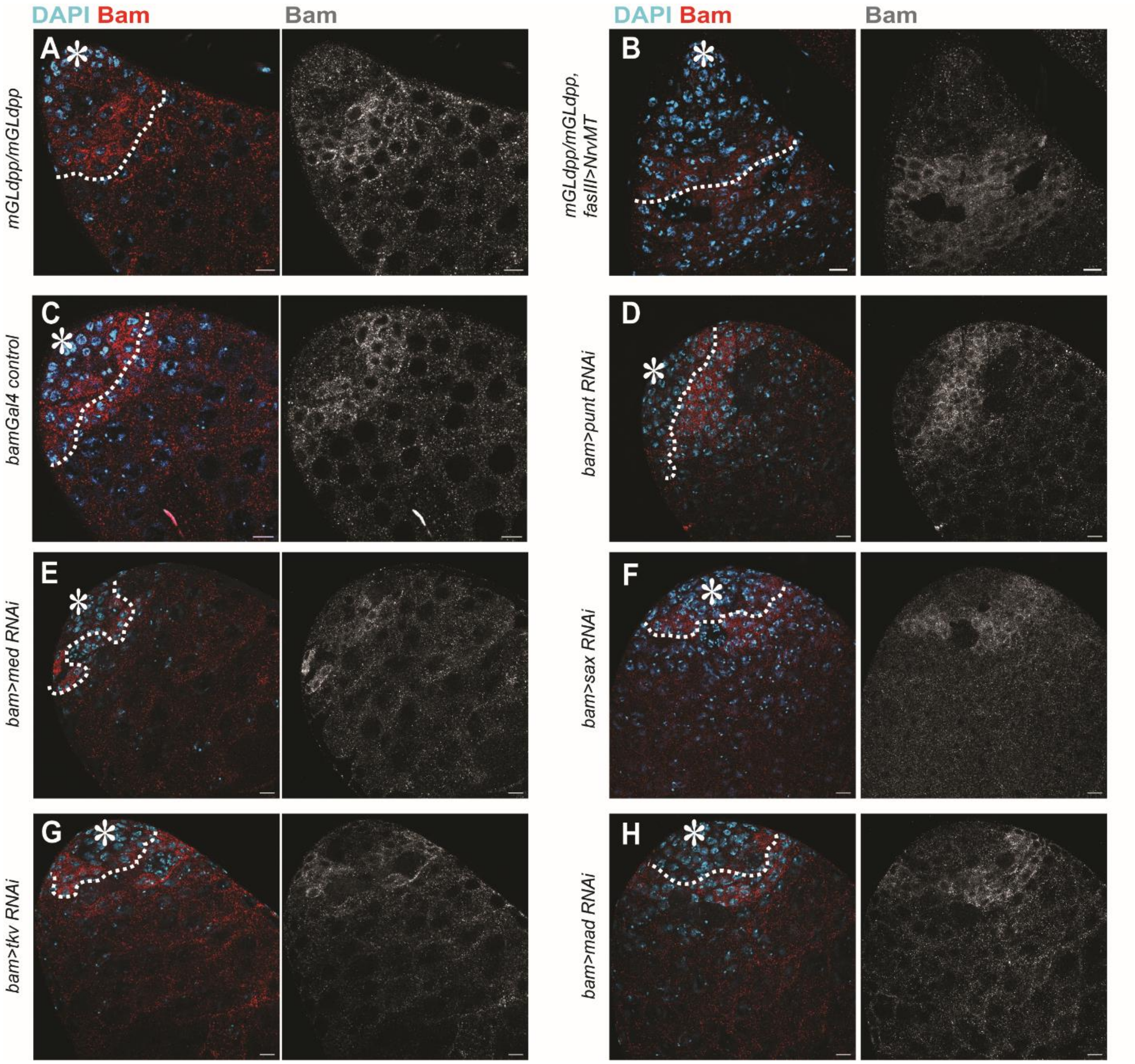


**Figure S6. BMP signal in SGs is required for timely upregulation of Bam.**

**A-H**) Representative Bam staining images of indicated genotypes. Boundary between 8-cell SGs and 16-cell SGs are divided by white broken lines. Asterisks indicate approximate location of the hub. All scale bars represent 10 μm. Fixed samples were used for all images.

### Legends for other supplemental materials

#### Video S1

A representative time-lapse movie of a testis tip (corresponding to Figure S4A). Time-interval: 10min. Scale bar: 10µm.

#### Video S2

A representative time-lapse movie of a testis tip (corresponding to Figure S4B). Time-interval: 10min. Scale bar: 10µm.

#### Source data

Numerical values of all graphs are provided in this excel spreadsheet.
